# A zebrafish model of COVID-19-associated cytokine storm syndrome reveals differential proinflammatory activities of Spike proteins of SARS-CoV-2 variants of concern

**DOI:** 10.1101/2021.12.05.471277

**Authors:** Sylwia D. Tyrkalska, Alicia Martínez-López, Ana B. Arroyo, Francisco J. Martínez-Morcillo, Sergio Candel, Diana García-Moreno, Pablo Mesa-del-Castillo, María L. Cayuela, Victoriano Mulero

## Abstract

The sudden and unexpected appearance of the COVID-19 pandemic turned the whole world upside down in a very short time. One of the main challenges faced has been to understand COVID-19 patient heterogeneity, as a minority develop life-threatening hyperinflammation, the so-called cytokine storm syndrome (CSS). Using the unique advantages of the zebrafish model we report here the proinflammatory role of Spike (S) proteins from different SARS-CoV-2 variants of concern after injection into the hindbrain ventricle, a cavity filled with cerebrospinal fluid to which immune cells can be easily recruited and that mimics the alveolar environment of the human lung. We found that wild type/Wuhan variant S1 (S1WT) protein promoted neutrophil and macrophage recruitment, local and systemic hyperinflammation, emergency myelopoiesis, and hemorrhages. In addition, S1γ protein was more proinflammatory and S1δ was less proinflammatory than S1WT and, strikingly, S1β promoted delayed and long-lasting inflammation. Pharmacological inhibition of the canonical inflammasome robustly alleviated S1 protein-induced inflammation and emergency myelopoiesis. In contrast, genetic inhibition of angiotensin-converting enzyme 2 strengthened the proinflammatory activity of S1, and the administration of angiopoietin (1-7) fully rescued S1-induced hyperinflammation and hemorrhages. These results shed light into the mechanisms orchestrating the COVID-19-associated CSS and the host immune response to different SARS-CoV-2 S protein variants.

**Highlights:**

- S proteins of SARS-CoV-2 promote hyperinflammation, neutrophilia, monocytosis and hemorrhages in zebrafish.
- S protein effects in zebrafish are mediated via the canonical inflammasome and the Ace2/Angiopoietin (1-7) axis.
- Delta S1 is less proinflammatory than wild type S1 and fails to induce emergency myelopoiesis in zebrafish.
- Naïve and primed human white blood cells are unable to respond to S proteins.

## INTRODUCTION

By the end of 2019, a new viral disease-causing pneumonia had been reported in Wuhan, China (Zhu et al., 2020). Metagenomic RNA sequencing revealed that a new betacoronavirus called Severe Acute Respiratory Distress Syndrome Coronavirus-2 (SARS-CoV-2) was responsible for the outbreaks in China and soon in the rest of the world. The disease was named Coronavirus disease 2019 (COVID-19) and was announced as the second pandemic of the twenty-first century by the World Health Organization (WHO) on March 11th 2020 (Cucinotta and Vanelli, 2020).

COVID-19 is believed to be transmitted via respiratory droplets and fomites. When the viral units are inhaled into the respiratory tract through the nasopharyngeal mucosal membranes, SARS-CoV-2 binds to the epithelial cells and starts replicating and migrating down to the airways reaching the alveolar epithelial cells in the lungs (Huang et al., 2020a). SARS-CoV-2 infection in humans varies and may manifest itself as mild symptoms to severe respiratory failure. The most common symptoms of COVID-19 include fever, dry cough, malaise, myalgia, vomiting, diarrhea, and abdominal pain (Guan et al., 2020; Huang et al., 2020a). Moreover, neurological, musculoskeletal and cardiovascular failures have lately been included in a list of potential COVID-19 complications as a result of multiple organ failure that can even lead to death (Wu and McGoogan, 2020). Although the clinical manifestations of COVID-19 differ with age, all ages of the population are susceptible to SARS-CoV-2 infection (Chen et al., 2020). The first signs of the disease become evident after an incubation period of 1–14 days, most commonly around day 5 (Wu and McGoogan, 2020).

The phylogenetic analysis of the whole genome of a novel SARS-CoV-2 virus shows close similarity with SARS-related coronaviruses (Viruses, 2020). Indeed, it has been shown that it shares 79% genome sequence identity with SARS-CoV and 50% with MERS-CoV (Lu et al., 2020). SARS-CoV-2 consists of a long single-stranded positive-sense RNA molecule, which is surrounded by a lipid envelope that anchors many structural viral glycoproteins (Heaton and Randall, 2011). The viral genome is organized in ten open reading frames (ORFs) encoding nonstructural polyproteins (67%) and accessory or structural proteins (33%) (Chan et al., 2020; Mittal et al., 2020). Four major structural proteins are encoded by SARS-CoV-2 genome: spike (S), envelope (E), membrane (M) and nucleocapsid (N) (Srinivasan et al., 2020). Spike protein has lately become the most studied structural protein among the viral proteins due to its role in viral recognition by the host and ability to mutate, thus generating new SARS-CoV-2 variants.

Spike protein is a trimeric glycoprotein that is encoded by ORF2 in the viral genome. It is formed by a membrane-distal S1 subunit and a membrane-proximal S2 subunit, which form homotrimers in the virus envelope (Yoshimoto, 2021). The Spike S1 subunit recognizes the host cellular receptor ACE2 via its receptor-binding domain (RBD) located at the C-terminal, while its N-terminal is composed of N-terminal domain (NTD) (Örd et al., 2020). The S2 subunit is essential for the membrane fusion that initiates virus entry (Walls et al., 2020). Recognition of the host cell receptor is followed by the structural rearrangements of SARS-CoV-2 S protein (Cai et al., 2020). More specifically, RBD domain, which is situated in the external subdomain, opens to bind to the ACE2 protein in the host cell. This results in S protein binding to two more ACE2 proteins to form the spike protein bound to three ACE2 proteins (Yoshimoto, 2021). Subsequently, the spike protein complex is cleaved by furin and TMPRSS2 to release the ACE2-S1 fragments (Bestle et al., 2020; Shang et al., 2020). Proteolytical cleavage can happen at two cleavage sites to separate S1 and S2 (Dai and Gao, 2021). As the S2 domain remains and is already primed for viral entry into the host cell, the refolding process of the spring-loaded S2 subunit initiates host and viral membranes fusion.

Viral genomes can undergo adaptive mutations that may provoke alterations in the virus’s pathogenic potential. It has been seen that even a single amino acid exchange can drastically affect and perturb the ability of a virus to evade the host immune system and hence complicate the progression of vaccine development against the virus. Unfortunately, SARS-CoV-2, like other RNA viruses, is prone to genetic evolution and develops mutations over time, resulting in the emergence of multiple variants that may have different characteristics compared to their ancestral strains (2021). Of the many different variants of SARS-CoV-2 that have evolved to date, this study concentrates on the three of most immediate concern and particular interest: Beta (lineage B.1.351), Gamma (lineage P.1) and Delta (lineage B.1.617.2) (Duong, 2021; Tao et al., 2021).

SARS-CoV-2 Beta variant was first detected in South Africa in October 2020, showing many new mutations especially in the region of S protein. It was observed that this variant spread more rapidly, and its prevalence was higher among young people with no underlying health conditions, and frequently resulted in serious illness. Moreover, Beta variant was able to attach to human cells more easily because of three mutations in the receptor-binding domain (RBD) within S protein: N501Y, K417N, and E484K (Duong, 2021). SARS-CoV-2 Gamma variant arose in Brazil in early 2021, showing 17 unique amino acid changes, ten of which were in its S protein, including the three mutations of most concern, N501Y, E484K and K417T, which change the stability of RBD-hACE2 complex affecting the binding affinity of RBD to hACE2 (Shahhosseini et al., 2021; Voloch et al., 2021). Gamma variant infections are 2.2 times more transmissible, have the same ability to infect both adults and older persons and can produce a ten-fold increase in viral load compared to persons infected by other variants (Duong, 2021). SARS-CoV-2 Delta variant was first discovered in India in late 2020, mutations in the S protein causing substitutions in T478K, L452R and P681R. These mutations are known to affect the transmissibility of the virus, and it is thought to be one of the most transmissible respiratory viruses known to date; and they are also involved in immunoescape (Duong, 2021; Mishra et al., 2021; Planas et al., 2021).

Nowadays, the zebrafish model is one of the fastest growing animal models used for reproducing all types of human diseases, such as cardiovascular diseases, infectious diseases, cancer, neurodegeneration diseases, hematopoietic disorders, and many more. Its small size, large number of eggs generated at a time and external fertilization, the transparency of the embryos, rapid development and easy maintenance in the laboratory make the zebrafish an attractive model for research in many fields (Diniz et al., 2015; Howe et al., 2013; Kimmel et al., 1995; Spence et al., 2008). Moreover, genetics and physiologic similarities with mammals increase their value as a powerful tool (MacRae and Peterson, 2015). Since the beginning of COVID-19 pandemic, many laboratories have started modelling the disease in zebrafish. Although, zebrafish does not have lungs, the presence of a hindbrain ventricle, a cavity filled with cerebrospinal fluid into which immune cells can be recruited, mimics the alveolar environment in the human lungs. This is a convenient injection site to study host and viral factors involved in local and whole-body immune response and innate immunity in general (Cambier et al., 2014; Herbomel et al., 1999; Torraca et al., 2015). Here, we describe a zebrafish model to study the immune response driven by different Spike variants of SARS-CoV-2 and explain the different proinflammatory activities that contribute to COVID-19-associated cytokine storm syndrome (CSS) (Fajgenbaum and June, 2020).

## RESULTS

### Wild type S1 is highly proinflammatory in zebrafish

Zebrafish are proving to be good models for several human diseases, including viral infections (Antoine et al., 2014; Espín-Palazón et al., 2016; Sullivan et al., 2021). Here we present a novel zebrafish model for COVID-19 using the recombinant S1 domain from wild type Spike protein (S1WT) injected into the hindbrain of 48-hour postfertilization (hpf) zebrafish larvae. The model does not need the virus to be used for the infection, which makes it easier, safer and cheaper to study the CSS associated to COVID-19. Injection of S1WT induced a robust recruitment of innate immune cells, namely neutrophils (*mpx:eGFP*) and macrophages (*mfap4:mcherry*), to the site of the injection. By six hours postinjection (hpi) both neutrophils and macrophages were present in the hindbrain of S1WT-injected larvae, in contrast to the control larvae injected with water or BSA, where the numbers of both cell types were insignificant (Figures 1A, 1B, S1A and S1B). This difference remained until 12 and 24 hpi, suggesting that S1WT can be recognized by the zebrafish immune system (Figures 1A and 1B). The same pattern can be seen by counting the number of neutrophils and macrophages in the head of the injected larvae (Figures 1A, 1B, S1A and S1B). Moreover, the total number of neutrophils and macrophages in the whole body was higher after S1WT injection compared with their control siblings at 24 hpi. This indicates that S1WT injected into the hindbrain was able to activate emergency hematopoiesis leading to increased production of neutrophils and macrophages.

**Figure 1:**
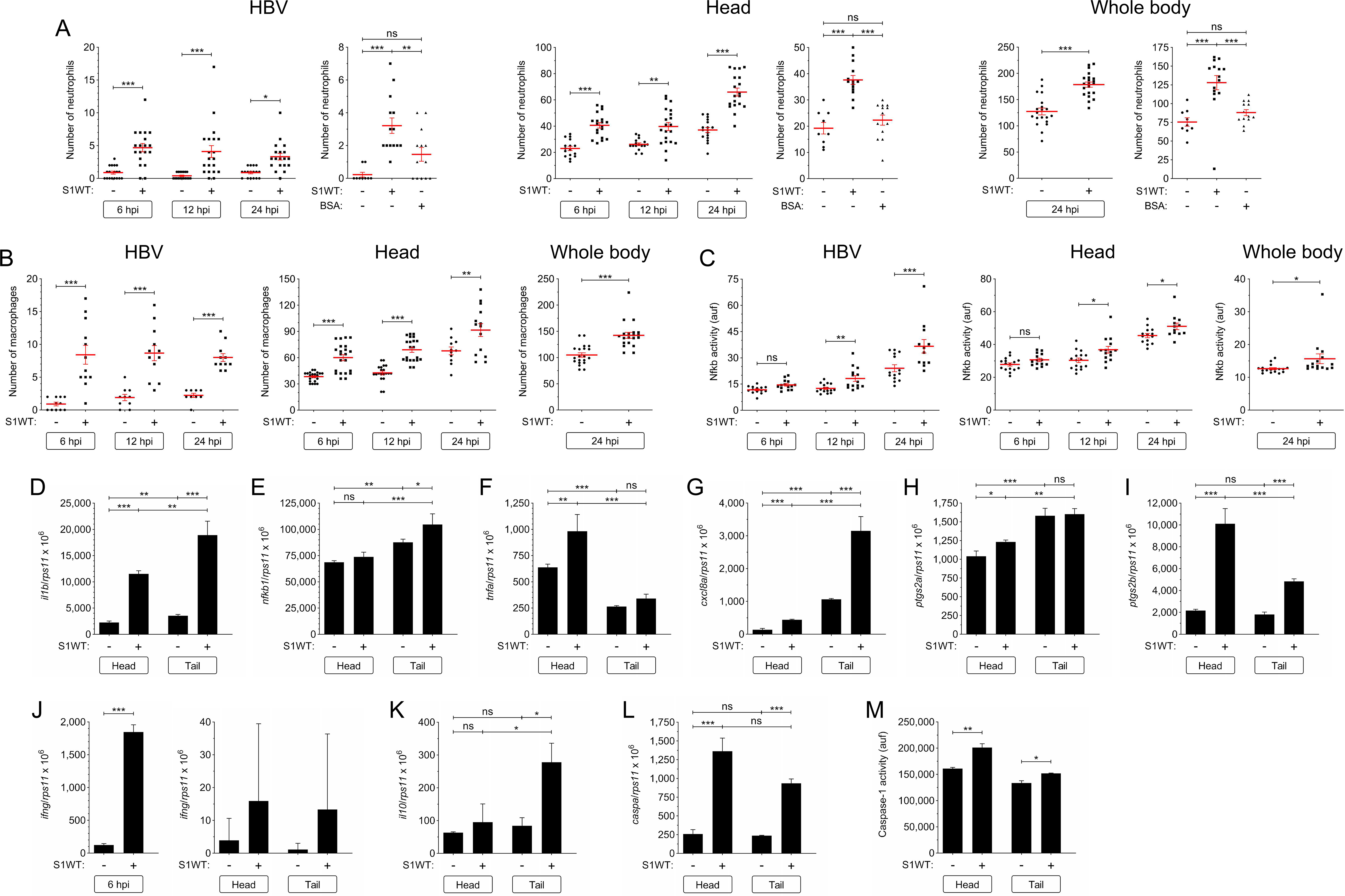
Wild type S1 is highly proinflammatory in zebrafish. Recombinant S1WT or BSA were injected in the hindbrain ventricle (HBV) of 2 dpf *Tg(mpx:eGFP)* (A), *Tg(mfap4:mCherry)* (B), *Tg(NFkB-RE:eGFP)* (C) and wild type (D-N) larvae. Neutrophil (A) and macrophage (B) recruitment and number, and Nfkb activation (C) were analyzed at 6, 12 and 24 hpi by fluorescence microscopy, the transcript levels of the indicated genes (D-L) were analyzed at 12 hpi by RT-qPCR in larval head and tail (except for *ifng* that was also analyzed at 6 hpi in the head), and caspase-1 activity was determined at 24 hpi using a fluorogenic substrate (M). Each dot represents one individual and the mean ± S.E.M. for each group is also shown. P values were calculated using one-way ANOVA and Tukey multiple range test. ns, not significant, *≤p0.05, **p≤0.01, ***p≤0.001. auf, arbitrary units of fluorescence.

To check how the inflammation process proceeds in the fish injected with S1WT, we first analyzed the expression pattern of the master regulator of the inflammatory response Nfkb using the reporter line *nfkb:eGFP*. Although at 6 hpi there was no increase in the fluorescent signal levels of Nfkb in the site of the injection or the head, increased fluorescence started to appear at 12 and continued until 24 hpi (Figures 1C and S1C). Nfkb had not only increased locally, but a slight increase was also seen globally in the whole body at 24 hpi (Figures 1C and S1C). These results suggest that S1WT is not only able to initiate the immune response locally at the site of the injection in zebrafish, but also to cause a systemic inflammation that spreads throughout the zebrafish body.

To confirm these results, we used RT-qPCR to measure the transcript levels of proinflammatory genes in the head (locally) and rest of the body (systemically) at 12 hpi. The transcript levels of *il1b*, *tnfa*, *cxcl8a*, *ptgs2a* and *ptgs2b* increased locally in the head, whereas *il1b*, *nfkb1*, *cxcl8a* and *ptgs2b* also increased in the rest of the body, further confirming the systemic inflammation triggered by S1WT (Figures 1D-1I). Curiously, the mRNA levels of the gene encoding interferon γ (*Ifng*) were seen to have drastically increased at 6 hpi but they returned to basal levels at 12 hpi (Figure 1J). The mRNA levels of the gene encoding the antiinflammatory cytokine Il10 increased systemically in the S1WT injected larvae (Figure 1K).

Recently, it has been shown that inflammasome plays an important role in the SARS-CoV-2 infection (Vora et al., 2021). To check whether the injection of S1WT into the hindbrain of the zebrafish larvae is able to activate the expression of genes other than proinflammatory genes, RT-qPCR was used to measure the transcript levels of Caspase a (Caspa, the functional homologue of human CASP1) The *caspa* transcript levels increased at both local and systemic levels (Figure 1L). Moreover, caspase-1 activity also increased in both the head and the rest of the body of S1WT-injected larvae (Figure 1M).

### Full-length S protein phenocopies the effects of S1 protein in zebrafish

To verify the results obtained with the zebrafish model for COVID-19 using recombinant S1WT protein injected into the hindbrain, we expressed wild type full-length S by forcing expression of its RNA, which was injected into the yolk sac of one-cell stage embryos. Using this strategy, full-length S was ubiquitously expressed as a membrane-anchored protein, as occurs in infected cells. Using this model, we observed Nfkb induction (Figure S2A) and higher transcript levels of *il1b*, *nfkb1*, *tnfa*, *cxcl8a*, *ptgs2a* and *ptgs2b, infg* and *il10* in the larvae expressing full-length S (Figures S2B-S2I) compared with control larvae. Although the mRNA levels of the genes encoding the most important inflammasome components Pycard (Apoptosis-associated speck-like protein containing a CARD, also known as Asc) and Caspa were not affected by the forced expression of full-length S (Figures S2J and S2K), caspase-1 activity was high in these fish compared with the controls (Figure S2L). Collectively, these results confirmed the strong proinflammatory effects of wild type S in zebrafish larvae.

### Recombinant S1, S1+S2 and E proteins are all proinflammatory in zebrafish

Initially, it was believed that only S protein, especially its S1 domain, was responsible for host immune system recognition, forming part of the outer sheath of the virus (Huang et al., 2020b). However, E protein, which is only believed to play a role in the viral assembly, also forms part of this sheath facing outwards, and is recognized by TLR2 to promote inflammation (Zheng et al., 2021). We therefore injected the hindbrain of zebrafish larvae with recombinant S1, S1+S2 or E proteins from the wild type variant and checked neutrophil and macrophage recruitment to the site of the injection 6, 12 and 24 hpi. The results showed that S1, S1+S2 and E were all able to recruit both type of immune cells to the site of the injection at similar levels (Figures S3A and S3B). Moreover, the number of neutrophils and macrophages in the head of the fish injected with S1, S1+S2 and E were similar, whereas the total number of neutrophils and macrophages in the whole body increased only in the larvae injected with either S1 or S1+S2 (Figures S3A and S3B). This suggests that S1 and S1+S2, but not E protein, were able to induce emergency myelopoiesis, as further confirmed by the ability of S1 and S1+S2 to induce *csf3a* transcript levels (Figure S3C). Very probably, the S1 domain was responsible for the induction of emergency myelopoiesis since the presence of S2 did not further increase it.

All viral proteins used in this experiment were able to increase Nfkb in hindbrain, head and whole body, all to a similar extent over the levels of their control siblings at all time points tested (Figure S3D). This was further confirmed by RT-qPCR experiments, whereby S1, S1+S2 and E were all able to induce similar transcript levels of *il1b*, *nfkb1*, *tnfa*, *cxcl8a*, *il10*, *infg*, *ptgs2a*, ptgs2b, *caspa* and *pycard* (Figures S3E-S3N). However, E protein systemically induced *il1b*, *cxcl8a*, *ptgs2a* and *ptgs2b* transcript levels, whereas S1+S2 induced more robustly those of *tnfa*, *infg* and *il10* (Figures S3E-S3N). Caspase-1 activity was increased by all viral proteins locally and systemically, but E protein in particular (Figure S3O).

### S1WT signals through the canonical inflammasome in zebrafish

The inflammasome also plays a pivotal role in COVID-19 (Vora et al., 2021). To ascertain whether this pathway also mediates the proinflammatory activity of S1WT in zebrafish, the larvae were treated with the specific caspase-1 inhibitor, VX-765, by bath immersion. Interestingly, although the pharmacological inhibition of caspase-1 failed to inhibit the recruitment of neutrophils to the hindbrain (Figure 2A), it was able to decrease the number of neutrophils in the head and in the whole body (Figure 2A). This result suggests that S1WT rapidly activated emergency myelopoiesis via the canonical inflammasome in zebrafish.

**Figure 2:**
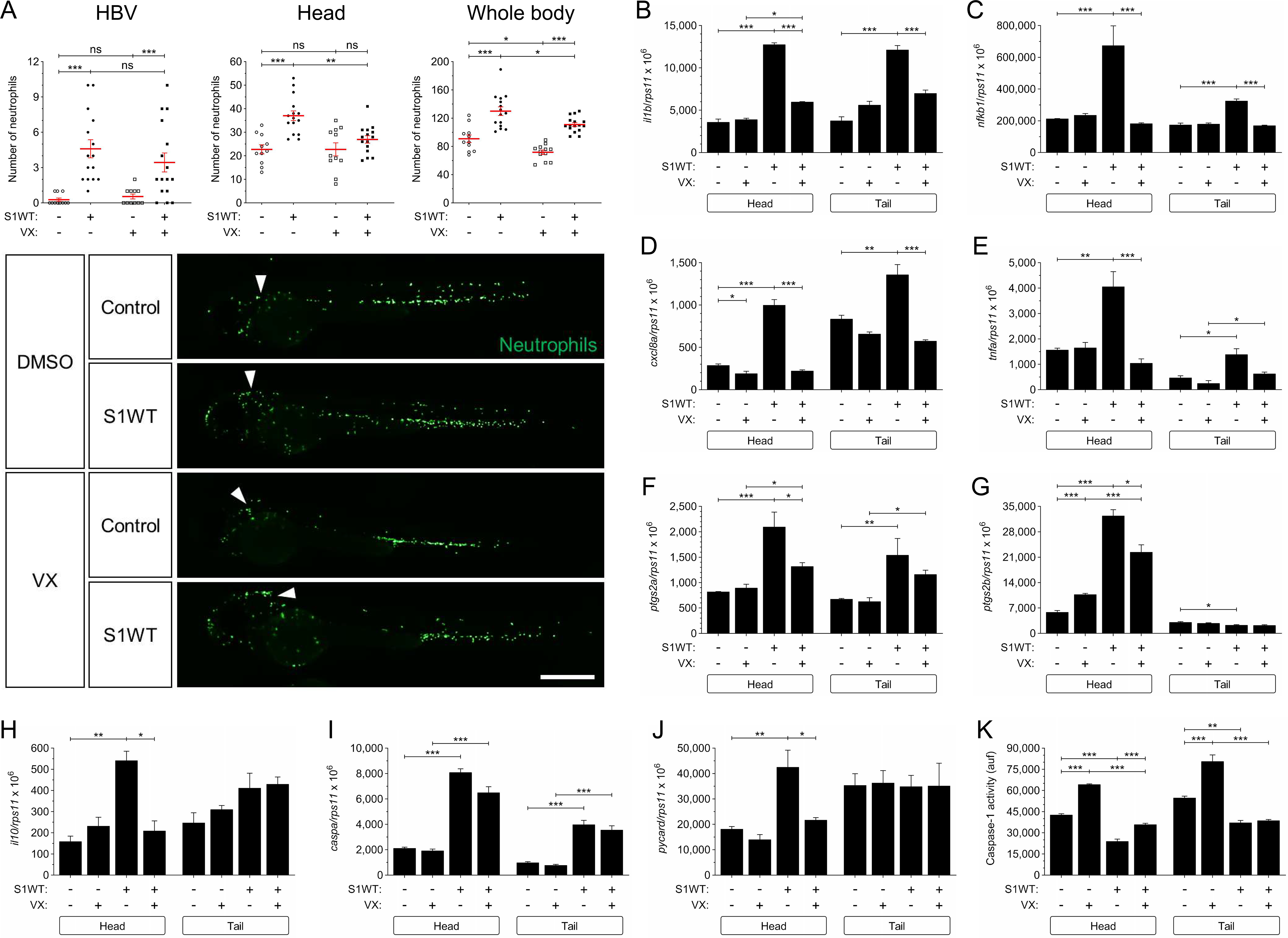
Wild type S1 signals through the canonical inflammasome in zebrafish. Recombinant S1WT was injected in the hindbrain ventricle (HBV, arrowheads) of 2 dpf *Tg(mpx:eGFP)* (A) and wild type (B-K) larvae in the presence of either DMSO or the caspase-1 inhibitor VX-765 (VX). Neutrophil recruitment and number were analyzed at 3 hpi by fluorescence microscopy (A), the transcript levels of the indicated genes were analyzed at 12 hpi by RT-qPCR (B-J), and caspase-1 activity was determined at 24 hpi using a fluorogenic substrate (K). Representative images of whole *Tg(mpx:eGFP)* larvae for each treatment are also shown (A). Each dot represents one individual and the mean ± S.E.M. for each group is also shown. P values were calculated using one-way ANOVA and Tukey multiple range test. ns, not significant, *≤p0.05, **p≤0.01, ***p≤0.001. Bar: 500 µm. auf, arbitrary units of fluorescence.

Next, we checked the transcript levels of proinflammatory genes in heads and the rest of the body of the fish injected with S1WT and treated with VX-765. It was found that pharmacological inhibition of the canonical inflammasome robustly decreased the induction of the proinflammatory genes caused by the injection of S1WT into the hindbrain of the zebrafish larvae (Figures 2B-2J). In addition, *il1b*, *nfkb1* and *cxcl8a* mRNA levels decreased at both local and systemic levels upon VX-765 treatment (Figure 2B-2D), whereas those of *tnfa*, *ptgs2a*, *ptgs2b* and *il10* decreased only at the injection site (Figure 2E-2H). Although no differences in *caspa* transcript levels were seen when S1WT injected larvae were treated with VX-765 (Figure 2I), those of *pycard* decreased to basal levels upon VX-765 treatment (Figure 2J). As expected, VX-765 decreased caspase-1 activity in the control and S1WT-injected larvae (Figure 2K). All these results indicate that SARS-CoV-2 S protein activates the canonical inflammasome in zebrafish.

### Ace2 deficiency exacerbates the proinflammatory activity of S1WT in zebrafish

Angiotensin-converting enzyme 2 (ACE2) is a zinc-containing metalloenzyme attached to the cell membrane, which is involved in the regulation of blood pressure through the hydrolysis of angiotensin (Ang)-II into Ang (1-7) (Bosso et al., 2020). Moreover, ACE2 serves as the entry point into the cells for different coronaviruses, among others SARS-CoV-2 (Gómez et al., 2020). Thus, we decided to check whether Ace2 was playing any role in the activation of zebrafish innate immunity mediated by recombinant S1WT protein. Surprisingly, although Ace2 deficiency (Figure S4) resulted in reduced neutrophil and macrophage recruitment to the injection site in both S1WT and flagellin injected larvae, the number of neutrophils or macrophages in the head and in whole larvae significantly increased in Ace2-deficient larvae at all time points (Figures 3A and 3B). Similarly, Nfkb activity was also higher in Ace2 deficient larvae injected with either S1WT or flagellin at the injection site, but also systemically, than in their control siblings (Figure 3C). It is worth mentioning that even the basal level of Nfkb in Ace2-deficient larvae injected with water were elevated in hindbrain, head and the whole body (Figure 3C).

**Figure 3:**
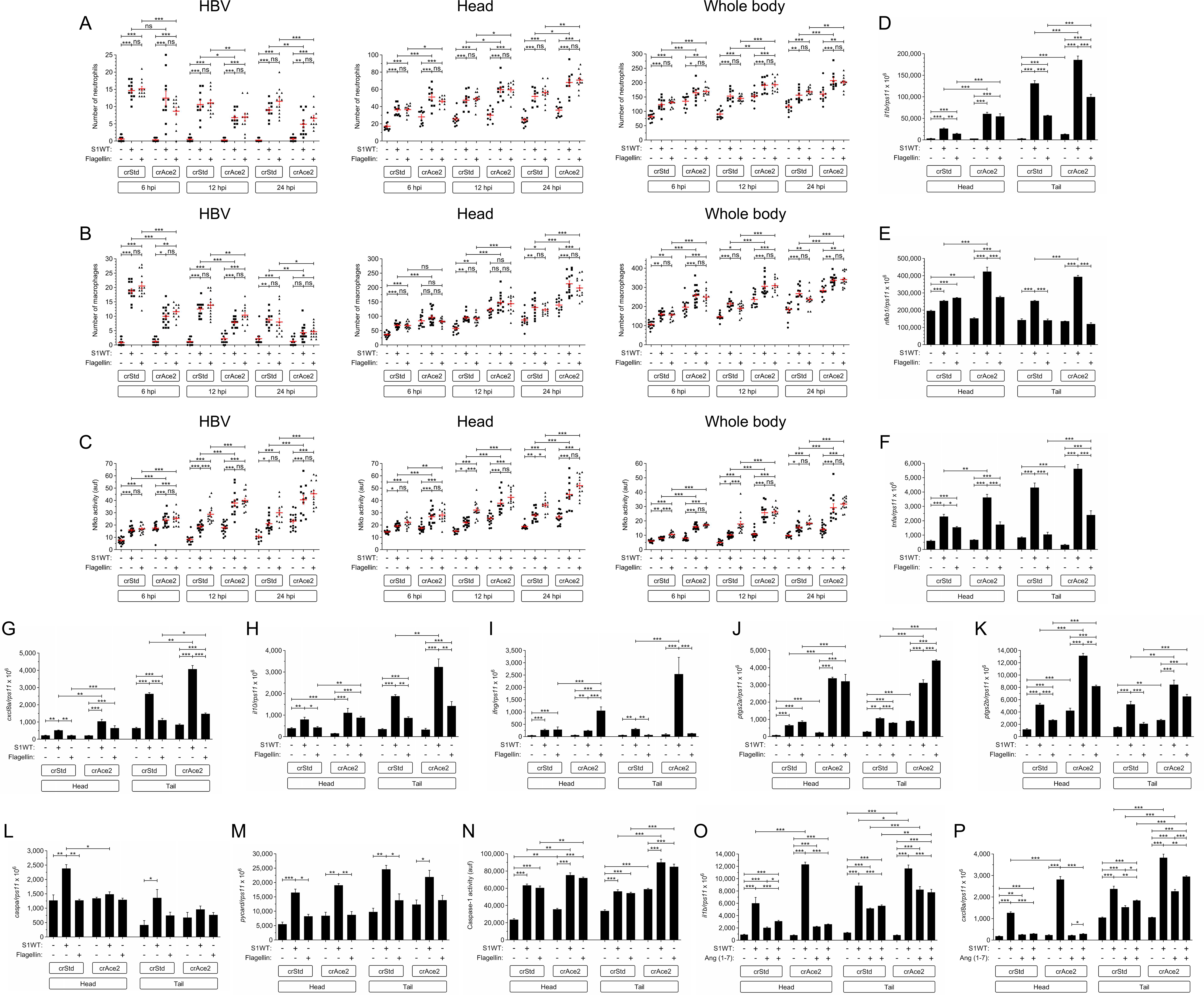
Ace2 deficiency exacerbates the proinflammatory activity of S1WT in zebrafish. One-cell stage zebrafish eggs of *Tg(mpx:eGFP)* (A), *Tg(mfap4:mCherry)* (B), *Tg(NFkB-RE:eGFP)* (C) and wild type (D-P) were microinjected with control or *ace2* crRNA/Cas9 complexes. At 2 dpf, recombinant S1WT or flagellin were injected alone or in combination with Ang (1-7) in the hindbrain ventricle (HBV) of control and Ace2-deficient larvae. Neutrophil (A) and macrophage (B) recruitment and number, and Nfkb activation (C) were analyzed at 6, 12 and 24 hpi by fluorescence microscopy, the transcript levels of the indicated genes were analyzed at 12 hpi by RT-qPCR (D-M, O, P), and caspase-1 activity was determined at 24 hpi using a fluorogenic substrate (N). Each dot represents one individual and the mean ± S.E.M. for each group is also shown. P values were calculated using one-way ANOVA and Tukey multiple range test. ns, not significant, *≤p0.05, **p≤0.01, ***p≤0.001. auf, arbitrary units of fluorescence.

The key anti-inflammatory role of Ace2 in zebrafish was further confirmed by the local and systemic increased transcript levels of *il1b*, nfkb1, *tnfa*, *cxcl8a*, *il10*, *infg*, *ptgs2a* and *ptgs2b* upon S1WT injection, and to some extent, flagellin, injection (Figures 3D-3K). Curiously, Ace2 deficiency failed to alter *caspa* and *pycard* mRNA levels (Figure 3L), and caspase-1 activity (Figure 3N) upon S1WT or flagellin injection. Strikingly, the injection of Ang (1-7) together with S1WT fully rescued the local hyperinflammation observed in Ace2-deficient larvae, as determined by quantitating the transcript levels of *il1b* and *cxcl8a* (Figures 3O and 3P).

### S1γ variant is more proinflammatory than the S1WT

The SARS-CoV-2 variant circulating in Brazil, called Gamma by the World Health Organization, is considered as a variant of concern. It has 17 amino acids substitutions, ten of which are present in the Spike protein (Faria et al., 2021). Hence, it was therefore checked *in vivo* whether there were any differences in the proinflammatory activities of recombinant S1WT and S1γ. It was found that fish injected with S1γ showed higher recruitment and a greater number of neutrophils and macrophages at all time points compared with the S1WT (Figures 4A and 4B). Similarly, the S1γ also induced higher Nfkb activity more rapidly (Figure 4C) and also, to some extent, proinflammatory gene transcripts (Figures 4D-4I) than S1WT. Notably, S1γ induced lower mRNA levels of *il10* both locally and systemically compared with S1WT (Figure 4J). Moreover, S1γ increased higher transcript levels of *caspa* (Figure 4K) and caspase-1 activity (Figure 4L). All these results suggest that S1γ is more proinflammatory and activates emergency myelopoiesis more robustly than S1WT.

**Figure 4:**
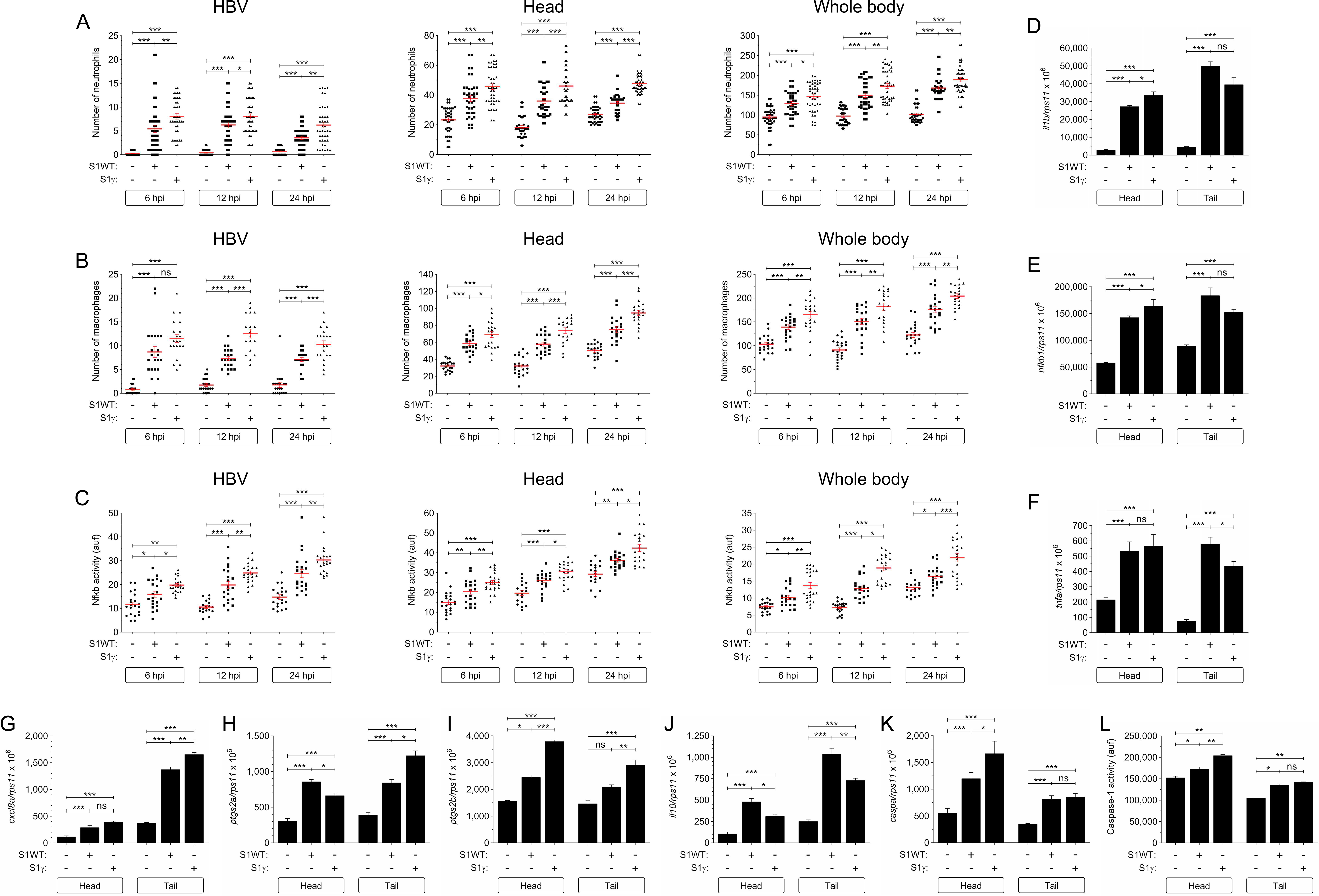
S1γ variant is more proinflammatory than the S1WT. Recombinant S1WT or S1γ were injected in the hindbrain ventricle (HBV) of 2 dpf larvae of *Tg(mpx:eGFP)* (A), *Tg(mfap4:mCherry)* (B), *Tg(NFkB-RE:eGFP)* (C) and wild type (D-L). Neutrophil (A) and macrophage (B) recruitment and number, and Nfkb activation (C) were analyzed at 6, 12 and 24 hpi by fluorescence microscopy, the transcript levels of the indicated genes were analyzed at 12 hpi by RT-qPCR (D-K), and caspase-1 activity was determined at 24 hpi using a fluorogenic substrate (L). Each dot represents one individual and the mean ± S.E.M. for each group is also shown. P values were calculated using one-way ANOVA and Tukey multiple range test. ns, not significant, *≤p0.05, **p≤0.01, ***p≤0.001. auf, arbitrary units of fluorescence.

### S1δ is less proinflammatory than the S1WT

Indian variant of SARS-COV2 also known as Delta variant, quickly spread around the world and is becoming the most dominant strain globally, is hence considered by the World Health Organization as a variant of concern. It possesses mutations is Spike protein that are known to affect transmissibility of the virus. Moreover, one of those mutations, L45R, is considered as the most interesting as it confers stronger affinity of the Spike protein for the ACE2 receptor and decreases the recognition capability of the host immune system (Starr et al., 2021; Zhang et al., 2021b). We found that the recruitment of neutrophils and macrophages to the injection site was more strongly attenuated in larvae injected with S1δ than with S1WT, while their total numbers were comparable with the control larvae injected with water (Figures 5A and 5B). Similarly, S1δ failed to induce Nfkb at any of the time points tested (Figure 5C) and was less potent at inducing the mRNA levels of inflammatory genes, except for *ptgs2b* (Figures 5D-5K). However, S1W and S1δ induced caspase-1 activity at similar levels (Figure 5L). Collectively, these results suggest that S1δ is much less proinflammatory than S1WT and fails to activate emergency hematopoiesis.

**Figure 5:**
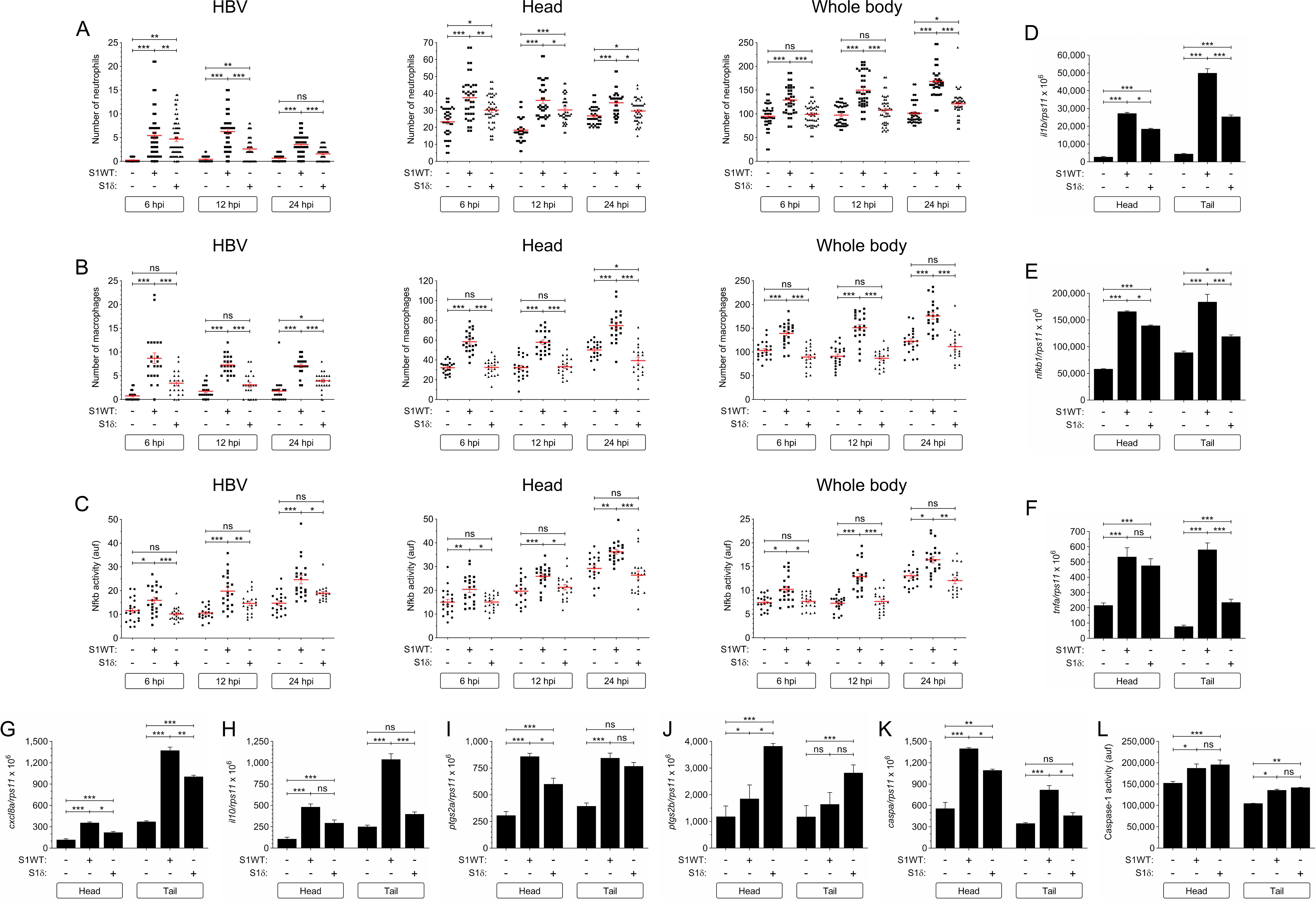
S1δ is less proinflammatory than the S1WT. Recombinant S1WT or S1δ were injected in the hindbrain ventricle (HBV) of 2 dpf larvae of *Tg(mpx:eGFP)* (A), *Tg(mfap4:mCherry)* (B), *Tg(NFkB-RE:eGFP)* (C) and wild type (D-L). Neutrophil (A) and macrophage (B) recruitment and number, and Nfkb activation (C) were analyzed at 6, 12 and 24 hpi by fluorescence microscopy, the transcript levels of the indicated genes were analyzed at 12 hpi by RT-qPCR (D-K), and caspase-1 activity was determined at 24 hpi using a fluorogenic substrate (L). Each dot represents one individual and the mean ± S.E.M. for each group is also shown. P values were calculated using one-way ANOVA and Tukey multiple range test. ns, not significant, *≤p0.05, **p≤0.01, ***p≤0.001. auf, arbitrary units of fluorescence.

### S1β shows delayed but stronger proinflammatory activity than the S1WT

South African variant of SARS-CoV-2 also known as Beta variant, has also been considered by the World Health Organization as a variant of concern. It possesses 8 mutations in Spike protein, especially within the receptor-binding domain (RBD), which makes it attach to human cells more easily and spread faster than other earlier variants of the virus (Tegally et al., 2021). It was observed that, although the initial influx (6 hpi) of neutrophils and macrophages to the injection site in response to S1β and S1WT was similar, or even slower for S1β, myeloid cell recruitment drastically increased at 12 and 24 hpi in larvae injected with S1β (Figures 6A and 6B). The same delayed induction pattern was observed in S1β-injected larvae for the Nfkb (Figure 6C), while the transcript levels of inflammatory genes were higher in S1β-injected larvae than their S1WT-injected counterparts, except for *tnfa* and *il10*, at 12 hpi (Figures 6D-6J). In addition, S1β induced higher *caspa* mRNA levels and caspase-1 activity in the injection site than S1WT (Figure 6K and 6L).

**Figure 6:**
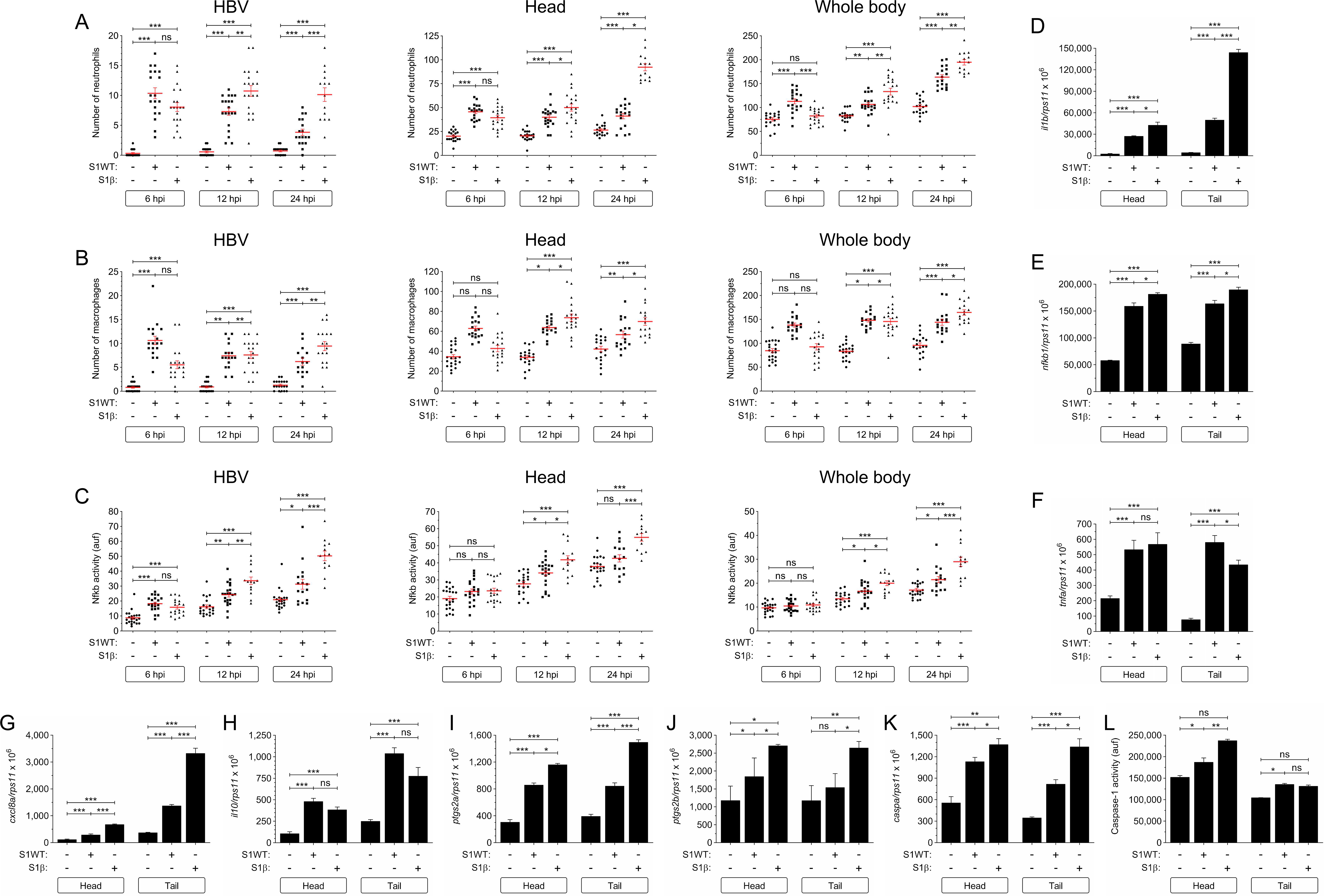
S1β shows delayed but stronger proinflammatory activity than the S1WT. Recombinant S1WT or S1β were injected in the hindbrain ventricle (HBV) of 2 dpf larvae of *Tg(mpx:eGFP)* (A), *Tg(mfap4:mCherry)* (B), *Tg(NFkB-RE:eGFP)* (C) and wild type (D-L). Neutrophil (A) and macrophage (B) recruitment and number, and Nfkb activation (C) were analyzed at 6, 12 and 24 hpi by fluorescence microscopy, the transcript levels of the indicated genes were analyzed at 12 hpi by RT-qPCR (D-K), and caspase-1 activity was determined at 24 hpi using a fluorogenic substrate (L). Each dot represents one individual and the mean ± S.E.M. for each group is also shown. P values were calculated using one-way ANOVA and Tukey multiple range test. ns, not significant, *≤p0.05, **p≤0.01, ***p≤0.001. auf, arbitrary units of fluorescence.

To further confirm that S1β was able to induce a sustained inflammatory response, *il1b* and *cxcl8a* mRNA levels were analyzed at 24 and 48 hpi. The results confirmed that, at 24 hpi, the larvae injected with S1β had higher *il1b* and *cxcl8a* levels than those injected with S1WT (Figure S5A). However, at 48 hpi the transcript levels of these genes had returned to basal levels in both S1WT and S1β (Figure S5B). These results show that S1β has delayed but longer lasting proinflammatory effects compared with the rest of S1 variants of concern (VOC).

### Recombinant S1 proteins provoke hemorrhages in zebrafish larvae

It has been noticed that patients with COVID-19 may present an increased risk of bleeding and develop different types of hemorrhages, mostly intracerebrally, with devastating consequences (Cheruiyot et al., 2021; Godier et al., 2021; Leasure et al., 2021). We, therefore, analyzed whether the injection of recombinant S1 proteins may produce hemorrhages by using a zebrafish line with labelled erythrocytes, i.e. *tg(gata1a:dsRed)*. Notably, the injection of S1 proteins into the hindbrain induced hemorrhages in the head of 25% of larvae and were already apparent at 6 hpi, remaining until 24 hpi (Figures 7A and 7B). Interestingly, all the S1 variants used gave rise to comparable percentages of larvae with hemorrhages (∼25%) (Figure 7C), while flagellin injection hardly induced hemorrhage (Figure 7C). As Ace2 and its catalytic product, Ang (1-7), were able to alleviate the proinflammatory effect of S1 in zebrafish larvae, we tested their impact on S1-induced hemorrhages. The results showed that although the genetic inhibition of Ace2 did not increase the percentage of larvae with hemorrhages triggered by S1WT, Ang (1-7) strongly decreased the percentage of larvae showing hemorrhages (Figure 7D). The ability of recombinant S1 proteins to give rise to hemorrhages in zebrafish rather than the injection itself was further confirmed by injecting full-length RNA of wild type S into one-cell stage embryos, leading to hemorrhages in the head of about 45% of the larvae at 48 hpf (Figures 7E and 7F).

**Figure 7:**
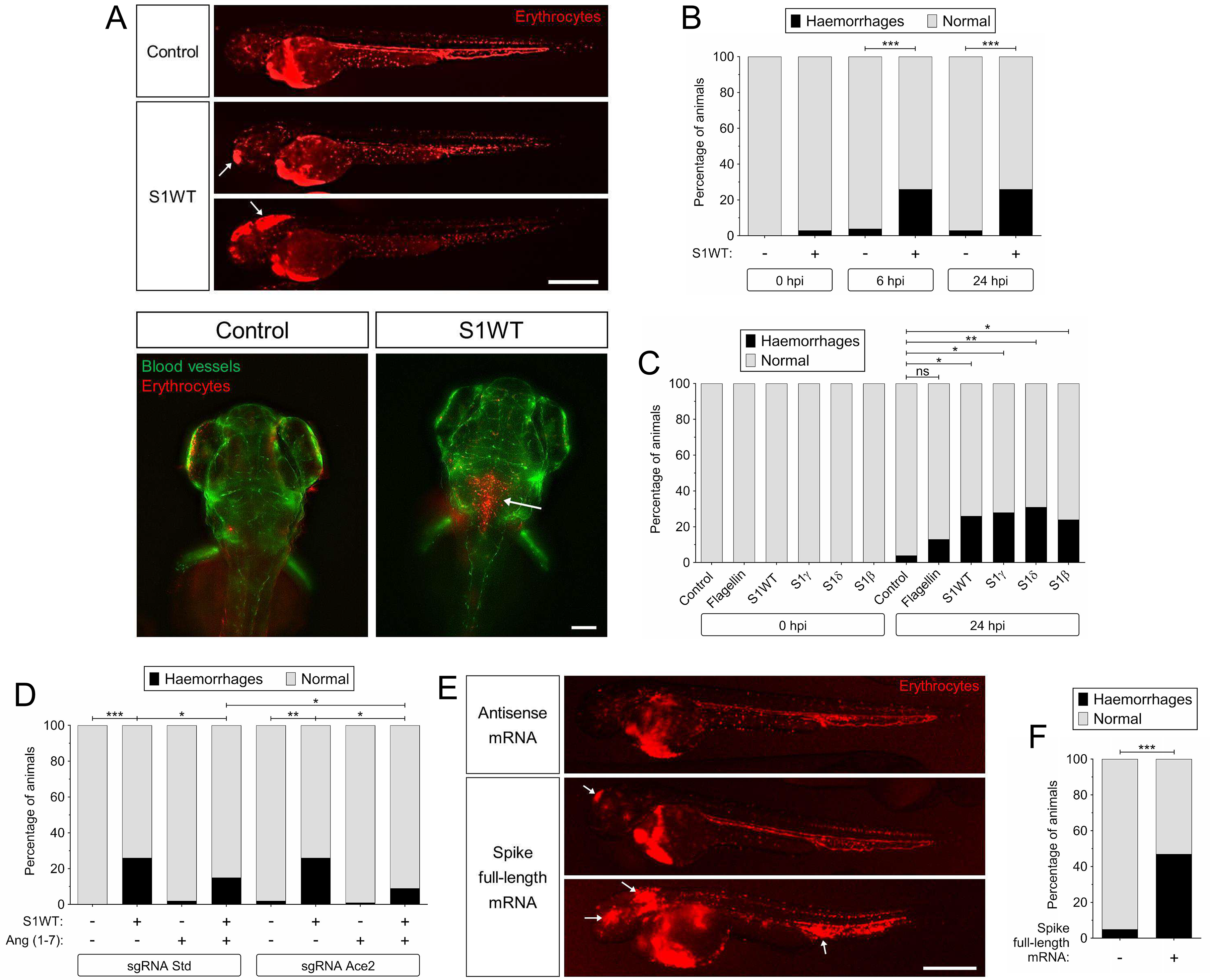
Recombinant S1 proteins provoke hemorrhages in zebrafish larvae. (A-D) One-cell stage zebrafish eggs of *Tg(gata1a:dsRed)*/*Tg(fli1a:eGFP)* were microinjected with control or *ace2* crRNA/Cas9 complexes. At 2 dpf, recombinant S1 proteins or flagellin were injected in the hindbrain ventricle (HBV) alone or combined with Ang (1-7). Hemorrhages were analyzed at 0, 6 and 24 hpi by fluorescence microscopy. (E,F) One-cell stage zebrafish eggs of *Tg(gata1a:dsRED)* were microinjected with control or full-length S RNAs and hemorrhages were analyzed at 48 hpf by fluorescence microscopy. The data are shown as the percentage of larvae showing hemorrhages. Representative images of the different experimental groups showing the hemorrhages (white arrows) are also shown (A, E). P values were calculated using Fisher’s exact test. ns, not significant, *p≤0.05, **p≤0.01, ***p≤0.001. Bars: 500 µm.

### Human mononuclear cells and neutrophils failed to respond to recombinant S proteins

The results obtained in zebrafish prompted us to analyze the impact of recombinant S1 proteins in human white blood cells, since controversial data have been reported in this respect. Unexpectedly, all recombinant S1 proteins tested failed to induce the transcript levels of genes encoding *TNFA*, *IL1B*, *PTGS2*, *CXCL8* and *ISG15* in peripheral blood mononuclear cells (PBMCs) (Figure 8A) and neutrophils (Figure 8B). We also used S1WT from another manufacturer, as well as S1+S2, and they also failed to stimulate PBMCs (Figure 8A) and peripheral blood neutrophils (Figure 8B). In contrast, recombinant E protein induced the same mRNA levels of all tested genes, except *ISG15*, with a similar potency as flagellin in both cell suspensions (Figures 8A and 8B). As it has been reported that human macrophages need to be primed to respond to recombinant S (Theobald et al., 2021), we next used white blood cells obtained from the synovial fluid of a patient with recent onset oligoarticular juvenile idiopathic arthritis. The results showed that although these cells were primed and flagellin and recombinant E protein induced higher transcript levels of *TNFA*, *IL1B*, *PTGS2* and *CXCL8* than in PBMCs and PBN, they were unable to respond to recombinant S proteins (Figure 8C).

**Figure 8.**
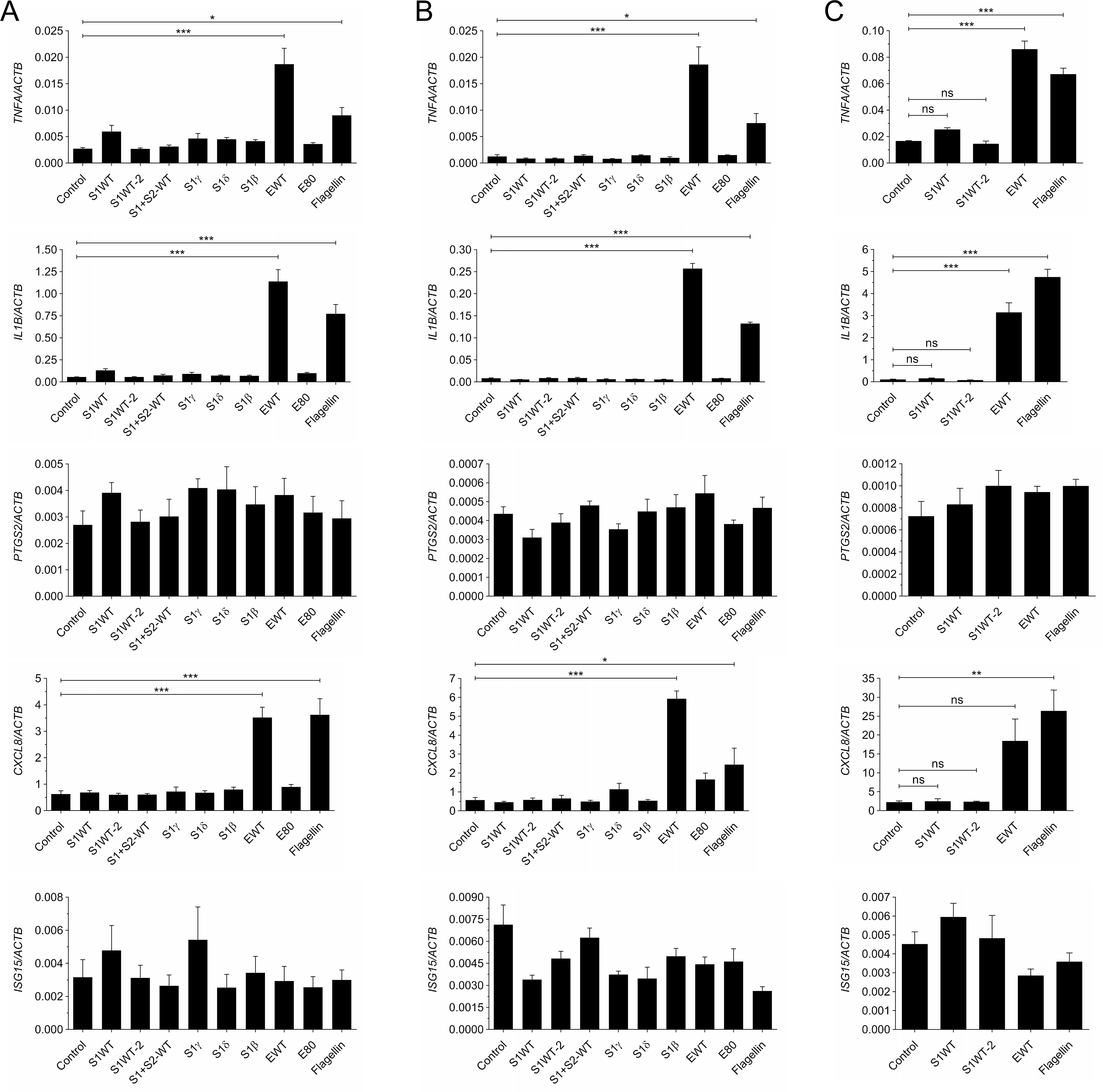
Human mononuclear cells and neutrophils failed to respond to recombinant S proteins. PBMCs (A) and neutrophils (B) from healthy donors, and white blood cells from the synovial fluid of a patient with recent onset oligoarticular juvenile idiopathic arthritis (C) were stimulated for 4 h with 1 µg/ml recombinant S1 (WT, γ, δ and β), S1+S2 (WT) and E (WT) proteins and the mRNA levels of the indicated genes analyzed by RT-qPCR. Data are shown as the mean ± S.E.M. of 3 technical replicates from 2 (A, B) or 1 (C) donors. P values were calculated using one-way ANOVA followed by Tukey multiple range test. ns, not significant, *≤p0.05, **p≤0.01, ***p≤0.001.

## DISCUSSION

The SARS-CoV-2 S protein is highly immunogenic and current COVID-19 vaccines are based on this property. However, there is controversy about the innate immune receptors and the signaling mechanisms involved in its recognition. While several studies have reported that S protein is able to interact and activate TLR4 in human THP-1 and HL-60 cells (Zhao et al., 2021) and murine macrophages and microglia (Olajide et al., 2021; Shirato and Kizaki, 2021; Zhao et al., 2021), others failed to demonstrate the activation of human or murine macrophages by S protein (Zheng et al., 2021). In addition, an elegant study has recently shown that S protein signals through TLR2 and activates the NLRP3 inflammasome in human macrophages obtained from COVID-19 convalescent patients, but not from healthy SARS-CoV-2 naïve individuals (Theobald et al., 2021). Although this study reveals that SARS-CoV-2 infection causes profound and long-lived reprogramming of macrophages resulting in augmented immunogenicity of the SARS-CoV-2 S-protein, it did not answer the question concerning the inability of naïve macrophages to respond to S proteins, despite expressing TLR2 which, however, was able to respond to zymosan (Theobald et al., 2021) and, intriguingly, to E protein (Zheng et al., 2021). We have also found that PBMCs and circulating neutrophils from healthy individuals failed to respond to S proteins, despite being able to recognize and mount an innate immune response to flagellin and E protein. Similarly, primed white blood cells, obtained from the synovial fluid of a patient with recent onset oligoarticular juvenile idiopathic arthritis, were also unable to respond to S protein, making these observations even more puzzling. In contrast, we found that zebrafish larvae were able to mount a quick innate immune response to recombinant S1 and S1+S2 obtained from different commercial sources, as well as to full-length S protein ubiquitously expressed in the larvae upon mRNA injection. Therefore, this model circumvents the limitations and discrepancies obtained with human and mouse macrophages concerning the immunogenic and proinflammatory properties of SARS-CoV-2 S protein and shed light into the COVID-19 associated CSS. Thus, we observed that S protein not only induced the local and systemic expression of genes encoding proinflammatory mediators, but also NFκB, neutrophilia and monocytosis. To the best of our knowledge, this is the first study showing that S protein is able to induce a dramatic neutrophilia and monocytosis, which are both involved in the pathogenesis of COVID-19 (Mei et al., 2020; Narasaraju et al., 2020).

Although the inflammasome has been found to participate in COVID-19, a mechanistic understanding of its involvement in COVID-19 progression remains unclear (Vora et al., 2021). Clinical trials with Anakinra, a modified human IL-1 receptor antagonist approved to treat rheumatoid arthritis, provided mixed results on patient benefits (Vora et al., 2021). We found that pharmacological inhibition of caspase-1, the effector of the canonical inflammasome, strongly alleviated the proinflammatory activity of S protein in zebrafish larvae. This effect was observed at two different levels: (i) the dampening of gene encoding proinflammatory mediators used in our study as a surrogate of the COVID-19-associated CSS, and (ii) the alleviation of neutrophilia. However, no differences in neutrophil recruitment to S protein was found between untreated and VX-765-treated larvae, indicating that the inflammasome was not involved in this process, as has been previously reported using a *Salmonella enterica* serovar Typhimurium infection model, where Cxcl8 and leukotriene B4 mediate neutrophil recruitment, while the inflammasome is responsible for the bacterial clearance (Tyrkalska et al., 2016). The impact of the inhibition of the inflammasome in both the S protein-induced inflammation and neutrophilia is an important observation, since dysregulated hematopoiesis has been observed in patients with severe COVID-19 (Wang et al., 2021), moreover the canonical inflammasome regulates the erythroid/myeloid decision of hematopoietic stem and progenitor cells (HSPCs) via cleavage of the master erythroid transcription factor GATA1 (Tyrkalska et al., 2019). Intriguingly, although E protein was also highly proinflammatory in zebrafish, it failed to promote emergency myelopoiesis, suggesting that E protein is unable to activate the inflammasome in HSPCs, despite its ability to activate the NLRP3 inflammasome in mouse macrophages (Zheng et al., 2021). Therefore, the zebrafish model developed here may contribute to understanding the contribution of the inflammasome to the CSS and the dysregulated hematopoiesis observed in severe COVID-19 patients, and to further understand the specific contribution of different viral structural proteins, as well as to identify therapeutic targets and novel drugs to treat COVID-19.

ACE2 is critical in the pathogenesis of COVID-19, not only as the viral receptor that allows the virus to invade the cells, but also as a major regulator of the renin angiotensin system (RAS)(Cousin et al., 2021). However, the role of ACE2 and RAS in COVID-19 is controversial. It has been hypothesized that viral infection promotes ACE2 internalization and downregulation following viral entry, promoting lower levels of Ang (1-7) but higher levels of Ang-II, which is linked to inflammation and fibrosis (Cousin et al., 2021). However, no significant changes in circulating RAS peptides or peptidases were found between uninfected and moderate COVID-19 patients (Files et al., 2021). We found that Ace2 inhibition exacerbated the proinflammatory effects of S protein in zebrafish larvae and, importantly, the administration of Ang (1-7) rescued the hyperinflammatory effects induced by S protein in Ace-2-deficient larvae. A similar effect of the Ace2/Ang (1-7) axis was observed in larvae injected with flagellin, suggesting a broad and strong anti-inflammatory effect of Ace2-derived Ang (1-7) rather than the neutralization of S protein by zebrafish Ace2. This is also supported by the inability of SARS-CoV-2 to replicate in zebrafish (Kraus et al., 2020; Laghi et al., 2021). Furthermore, although Ace2 deficiency had no impact on S1-induced hemorrhages, the administration of Ang (1-7) was able to reduce the percentage of S1-injected larvae showing hemorrhages. These results are of clinical relevance, since SARS-CoV-2 infection also induces vascular complications (Acharya et al., 2021) and Ang (1-7) has been found to be an useful therapeutic target for the treatment of cardiovascular disease, especially in patients with overactive RAS (Jiang et al., 2014). Our results point to a greater relevance of the ACE2/Ang (1-7)/Mas receptor axis than of the ACE/Ang-II/Angiotensin Type 1 receptor axis on COVID-19-associated CSS and vascular complications, and to the therapeutic potential of Ang (1-7) in this disease.

One of the most interesting observations of our study is the dramatic differences observed among the VOCs used. S1γ was found to be much more proinflammatory than S1WT, whereas S1δ showed an opposite behavior. Notably, S1δ failed to induce emergency myelopoiesis, despite being able to induce caspase-1 activity at similar levels as S1WT. As the inhibition of caspase-1 attenuated S1WT-induced emergency myelopoiesis, other mechanisms may be involved, such as the regulation of myeloid colony-stimulating factors by S proteins. Another possibility is that S1δ activates an inflammasome not involved in the regulation of hematopoiesis. Whatever the outcome, it is tempting to speculate that the reduced proinflammatory activity of S1δ reported here for the first time would facilitate high viral loads (∼1000 times higher) to be reached in the early stages of infection (Li et al., 2021), together with its enhanced infectivity via cell surface entry facilitated by its increased efficiency to cleave the full-length S to S1 and S2 (Liu et al., 2021) and to fuse membranes at low levels of cellular receptor ACE2 (Zhang et al., 2021a). As regards the S1β, we unexpectedly observed a delayed, but long-lasting, proinflammatory activity. Although the relevance of this observation in the contagiousness and pathogenesis of the SARS-CoV-2 Beta VOC remains to be determined, it has been reported that both Alpha and Beta VOCs produce tissue-specific cytokine signatures pathogenic patterns distinct from early strains in K18-hACE2 transgenic mice models (Radvak et al., 2021).

In summary, our zebrafish model developed to study COVID-19-associated CSS and the impact of different VOCs of SARS-CoV-2 S proteins, suggests that the canonical inflammasome and the ACE2/Ang (1-7) axis are key signaling pathways involved in the recognition of S protein by the host . In addition, the lower proinflammatory activity of S1δ may explain the high viral load reached by this VOC of SARS-CoV-2 in the early stages of infection. This model, therefore, is an excellent platform for the chemical screening of anti-inflammatory compounds to alleviate COVID-19-associated CSS.

## MATERIALS AND METHODS

### Animals

Zebrafish (*Danio rerio* H.) were obtained from the Zebrafish International Resource Center and mated, staged, raised and processed as described (Westerfield, 2000). The lines *Tg(mpx:eGFP)^i114^* (Renshaw et al., 2006), *Tg(mfap4:mCherry-F)^ump6^* (Phan et al., 2018), *Tg(NFkB-RE:eGFP)^sh235^* referred to as *nfkb:eGFP* (Kanther et al., 2011), *Tg(gata1a:DsRed)^sd2^* (Traver et al., 2003), *Tg(fli1:EGFP)^y^*^1^ (Delov et al., 2014) and casper (*mitfa^w2/w2^; mpv17^a9/a9^)* (White et al., 2008) were previously described. The experiments performed comply with the Guidelines of the European Union Council (Directive 2010/63/EU) and the Spanish RD 53/2013. The experiments and procedures were performed approved by the Bioethical Committees of the University of Murcia (approval number #395/2017).

### CRISPR, RNA and recombinant protein injections in zebrafish

crRNA for zebrafish *ace2* and tracrRNA were resuspended in Nuclease-Free Duplex Buffer to 100 µM. 1µl of each was mixed and incubated for 5 min at 95 ^0^C for duplexing. After removing from the heat and cooling to room temperature, 1.43 µl of Nuclease-Free Duplex Buffer was added to the duplex, giving a final concentration of 1000 ng/µl. Finally, the injection mix was prepared by mixing 1 µl of duplex, 2.55 µl of Nuclease-Free Duplex Buffer, 0.25 µl Cas9 Nuclease V3 (IDT, 1081058) and 0.25 µl of phenol red, giving final concentrations of 250 ng/µl of gRNA duplex and 500 ng/µl of Cas9. The prepared mix was microinjected into the yolk sac of one-to eight-cell-stage embryos using a microinjector (Narishige) (0.5–1 nl per embryo). The same amounts of gRNA were used in all the experimental groups. The efficiency of gRNA was checked by amplifying the target sequence with a specific pair of primers (Table 1) and the TIDE webtool (https://tide.nki.nl/).

*In vitro*-transcribed RNA was obtained following manufacturer’s instructions (mMESSAGE mMACHINE kit, Ambion). RNA was mixed in microinjection buffer and microinjected into the yolk sac of one-cell-stage embryos using a microinjector (Narishige; 0.5– 1 nl per embryo). The same amount of RNA was used for all the experimental groups.

Recombinant His-tagged Spike S1 wild type (#40591-V08B1), S1 (L18F, D80A, D215G, LAL242-244 deletion, R246I, K417N, E484K, N501Y, D614G) (variant β, cat. #40591-V08H15), S1 (L18F, T20N, P26S, D138Y, R190S, K417T, E484K, N501Y, D614G, H655Y) (variant γ, cat. #40591-V08H14) and (E154K, L452R, E484Q, D614G, P681R) (variant δ, cat. #40591-V08H19) (from Sino Biological); wild type Spike S1 (cat. #RP01262), Spike S1+S2 (cat. #RP01283LQ) and Envelope protein (E, cat. #RP01263) (from ABclonal), flagellin (Invivogen) or BSA (Sigma-Aldrich) at a concentration of 0.25 mg/ml supplemented with phenol red were injected into the hindbrain (1 nl).

### Chemical treatments

Two dpf larvae were manually dechorionated and treated for 1 or 4 h at 28 ^0^C by bath immersion with the caspase-1 inhibitor VX-765 (Belnacasan, Selleckchem) at a final concentration of 100 µM diluted in egg water supplemented with 0.1% DMSO. In some experiments, 24 hpf embryos were treated with 0.3% N-Phenylthiourea (PTU) to inhibit melanogenesis.

### Analysis of gene expression

Total RNA was extracted from whole larvae, head/tail larvae or human cell pellets with TRIzol reagent (Invitrogen) following the manufacturer’s instructions and treated with DNase I, amplification grade (1 U/mg RNA: Invitrogen). SuperScript IV RNase H Reverse Transcriptase (Invitrogen) was used to synthesize first-strand cDNA with random primer from 1µg of total RNA at 50 °C for 50 min. Real-time PCR was performed with an ABIPRISM 7500 instrument (Applied Biosystems) using SYBR Green PCR Core Reagents (Applied Biosystems). Reaction mixtures were incubated for 10 min at 95 °C, followed by 40 cycles of 15 s at 95 °C, 1 min at 60 °C, and finally 15 s at 95 °C, 1 min 60 °C, and 15 s at 95 °C. For each mRNA, gene expression was normalized to the *rps11* (zebrafish) or *ACTB* (human) content in each sample, using the Pfaffl method (Pfaffl, 2001). The primers used are shown in Table S1. In all cases, each PCR was performed with triplicate samples and repeated at least with two independent samples.

### Caspase-1 activity assays

The caspase-1 activity was determined with the fluorometric substrate Z-YVAD 7-Amido-4-trifluoromethylcoumarin (Z-YVAD-AFC, caspase-1 substrate VI, Calbiochem), as described previously (Angosto et al., 2012; Lopez-Castejon et al., 2008). In brief, 25–35 larvae were lysed in hypotonic cell lysis buffer (25 mM 4-(2-hydroxyethyl) piperazine-1-ethanesulfonic acid, 5 mM ethylene glycol-bis(2-aminoethylether)-N,N,Ń,Ń-tetraacetic acid, 5 mM dithiothreitol, 1:20 protease inhibitor cocktail (Sigma-Aldrich), pH 7.5) on ice for 10 min. For each reaction, 100 µg protein were incubated for 90 min at room temperature with 50 mM YVAD-AFC and 50 µl of reaction buffer (0.2% 3-[(3-cholamidopropyl)dimethylammonio]-1-propanesulfonate (CHAPS), 0.2 M 4-(2-hydroxyethyl) piperazine-1-ethanesulfonicacid, 20% sucrose, 29 mM dithiothreitol, pH 7.5). After incubation, the fluorescence of the AFC released from the Z-YVAD-AFC substrate was measured with a FLUOstart spectofluorometer (BGM, LabTechnologies) at an excitation wavelength of 405 nm and an emission wavelength of 492 nm. A representative caspase-1 activity graph out of three repeats is shown in figures.

### Imaging of zebrafish larvae

To study immune cell recruitment to the injection site and Nfkb activation, 2 dpf *mpx:eGFP*, *mfap4:mcherry* or *nfkb:egfp* larvae were anaesthetized in embryo medium with 0.16 mg/ml tricaine. Images of the hindbrain, head or the whole-body area were taken 3, 6, 12 and 24 h post-injection (hpi) using a Leica MZ16F fluorescence stereomicroscope. The number of neutrophils or macrophages was determined by counting visually and the fluorescence intensity was obtained and analyzed with ImageJ (FIJI) software (Schindelin et al., 2012).

*fli1a:EGFP;gata1a:dsRED* larvae previously treated with PTU were mounted for imaging in 1.6% low melting agarose in embryo medium with tricaine. Confocal images were acquired 24 h after hindbrain injection with the different recombinant S1 proteins on a widefield microscope (Leica THUNDER imager) with a 10x/0.32 objective using a DFC 9000 GTC sCMOS camera (Leica, Weztlar). The images are a maximum projection of a z-stack of 27 planes spaced at 3.8 microns and were processed with the Leica LVCC (Large Volume Computational Clearing) algorithm, which consists of a regularized Richardson-Lucy deconvolution preceded by a proprietary background subtraction (Leica ICC) that improves the deconvolution results on thick samples.

### Human PBMC, neutrophils and synovial fluid cell culture and treatments

All the experiments and procedures were performed as approved by the Ethical Clinical Research Committee of The University Hospital Virgen de la Arrixaca (approval number #5/2021). Peripheral blood was obtained from healthy donors and diluted 1/2 with physiological serum. The diluted blood was added to Lymphoprep (Alere Technologies AS, cat. #1114545) avoiding phase mixing. Subsequently, the tube was centrifuged for 30 min at 400x*g* without brake. PBMCs were collected from the white phase and neutrophils from the pellet of the sample. PBMC were then washed once with RPMI medium supplemented with 1% Penicillin/Streptomycin, 1% Glutamine and 10% FBS, and centrifuged for 5 min at 400x*g*. Finally, cells were resuspended in full RPMI medium to reach a final concentration of 10^6^ cells/ml. Neutrophils were washed with ACK lysis buffer (cat. #0000944064) to eliminate the erythrocytes and then twice with PBS. The remaining cells were resuspended in full RPMI medium to a final concentration of 10^6^ cells/ml. The synovial fluid -obtained from a 13-year-old female with recent onset of oligoarticular juvenile idiopathic arthritis -was composed of 87% mononuclear cells and 13% neutrophils. The fluid was treated with hyaluronidase for 10 min at room temperature and then centrifuged for 10 min at 400x*g*. The cell pellet was washed with ACK lysis buffer and then resuspended in supplemented RPMI medium to a final concentration of 10^6^ cells/ml. Cells were seeded at approximately 4×10^5^ cells per well in a 24-well plate and stimulated with 1µg/ml recombinant S1, S1+S2 and E proteins for 4 h at 37 °C. As controls, flagellin and the E protein pre-heated at 80 °C for 30 min were used at the same concentration.

### Statistical analysis

Data are shown as mean ± s.e.m., and the statistical differences among groups were determined by an analysis of variance and a Tukey multiple range test were. The differences between two samples were analyzed by Student’s *t*-test. A log-rank test was used to calculate the statistical differences in the survival of the different experimental groups.

## ACKNOWLEDGMENTS

We warmly thank I. Fuentes and P. Martínez for their excellent technical assistance and the staff of the Rheumatology Service of the University Hospital Virgen de la Arrixaca for collecting patient samples. We also thank Profs. S.A. Renshaw, P. Crosier, A.H. Meijer, D. Tobin and L.I. Zon for the zebrafish lines.

## Author contributions

VM and MLC conceived the study; SDT, AML, ABA, FJMM, SC and DGM performed the research; SDT, AML, ABA, FJMM, SC, DGM, PMdC, MLC and VM analyzed the data; and SDT and VM wrote the manuscript with minor contributions from other authors.

---

**Figure S1 (related to Figure 1).**
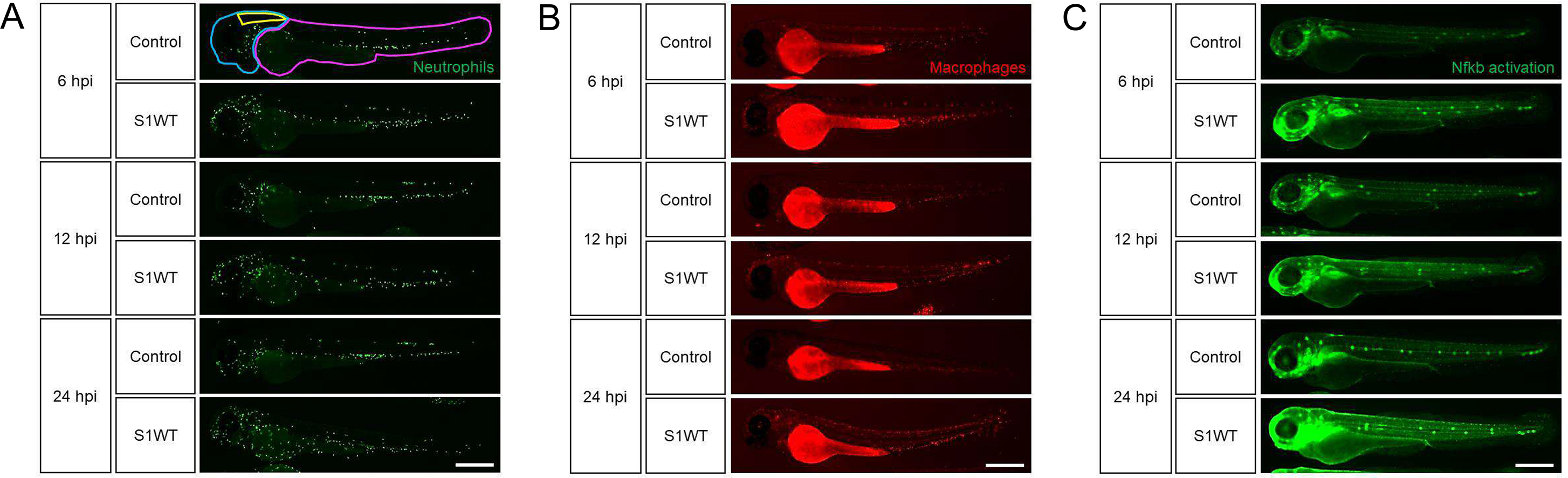
Wild type S1 is highly proinflammatory in zebrafish. Representative green and red fluorescence images of larvae of the different groups shown in Figures 1A-1C. Recombinant S1WT was injected in the hindbrain ventricle (HBV) of 2 dpf *Tg(mpx:eGFP)* (A), *Tg(mfap4:mCherry)* (B) and *Tg(NFkB-RE:eGFP)* (C) and neutrophils (A), macrophages (B) and Nfkb activation (C) visualized at 6, 12 and 24 hpi by fluorescence microscopy. The regions of interest used for quantitation in all experiments are indicated in A: hindbrain (yellow), head (blue) and rest of the body (violet). Bars: 500 µm.

**Figure S2 (related to Figure 1).**
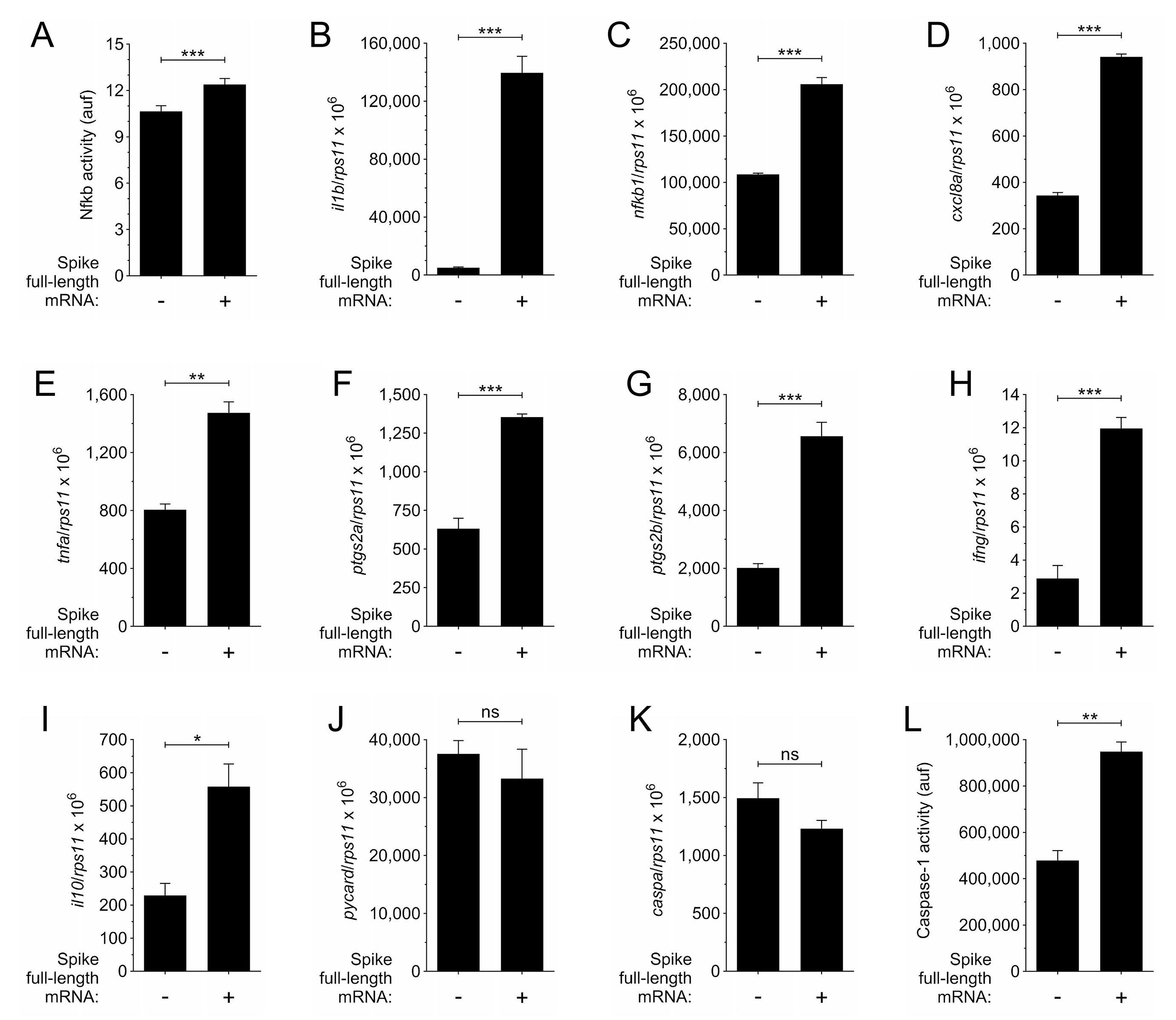
Full-length S protein phenocopies the effects of recombinant S1 protein in zebrafish. One-cell stage zebrafish eggs of *Tg(NFkB-RE:eGFP)* (A) or wild type (B-L) were microinjected with control or full-length S RNAs. Nfkb activation was analyzed at 48 hpf by fluorescence microscopy (A), the transcript levels of the indicated genes were analyzed at 48 hpf by RT-qPCR (B-K), and caspase-1 activity was determined at 72 hpf using a fluorogenic substrate (L). Data are shown as the mean ± S.E.M. obtained from 3 replicates. P values were calculated using one-way ANOVA followed by Tukey multiple range test. ns, not significant, *≤p0.05, **p≤0.01, ***p≤0.001.

**Figure S3 (related to Figure 1).**
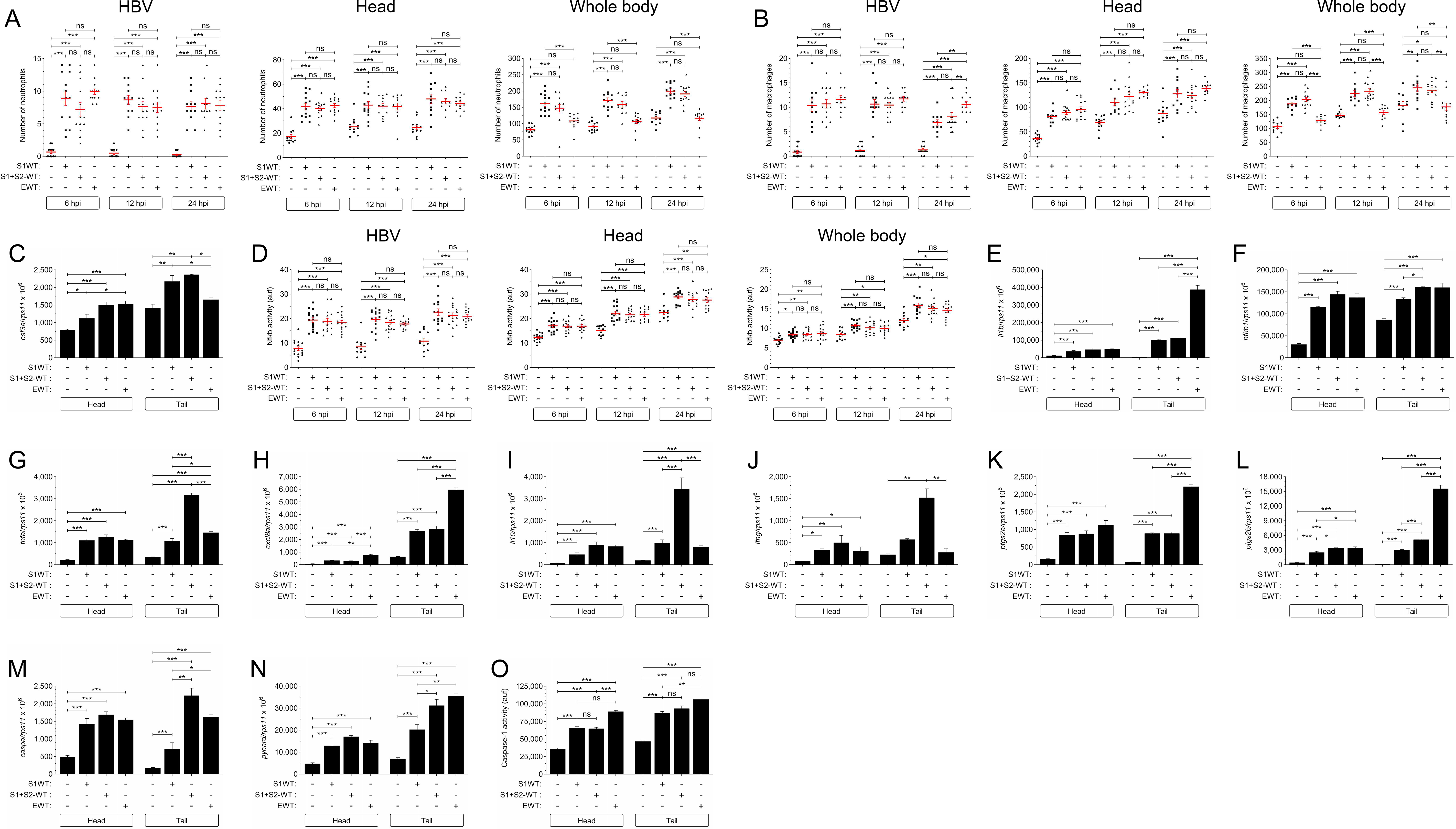
Recombinant S1, S1+S2 and E proteins are all proinflammatory in zebrafish. Recombinant S1, S1+S2 or E proteins from wild type SARS-CoV-2 were injected in the hindbrain ventricle (HBV) of 2 dpf larvae of *Tg(mpx:eGFP)* (A), *Tg(mfap4:mCherry)* (B), *Tg(NFkB-RE:eGFP)* (D) and wild type (C, E-O). Neutrophil (A) and macrophage (B) recruitment and number, and Nfkb activation (D) were analyzed at 6, 12 and 24 hpi by fluorescence microscopy, the transcript levels of the indicated genes were analyzed at 12 hpi by RT-qPCR (C, E-N), and caspase-1 activity was determined at 24 hpi using a fluorogenic substrate (O). Each dot represents one individual and the mean ± S.E.M. for each group is also shown. P values were calculated using one-way ANOVA and Tukey multiple range test. ns, not significant, *≤p0.05, **p≤0.01, ***p≤0.001.

**Figure S4 (related to Figure 3).**
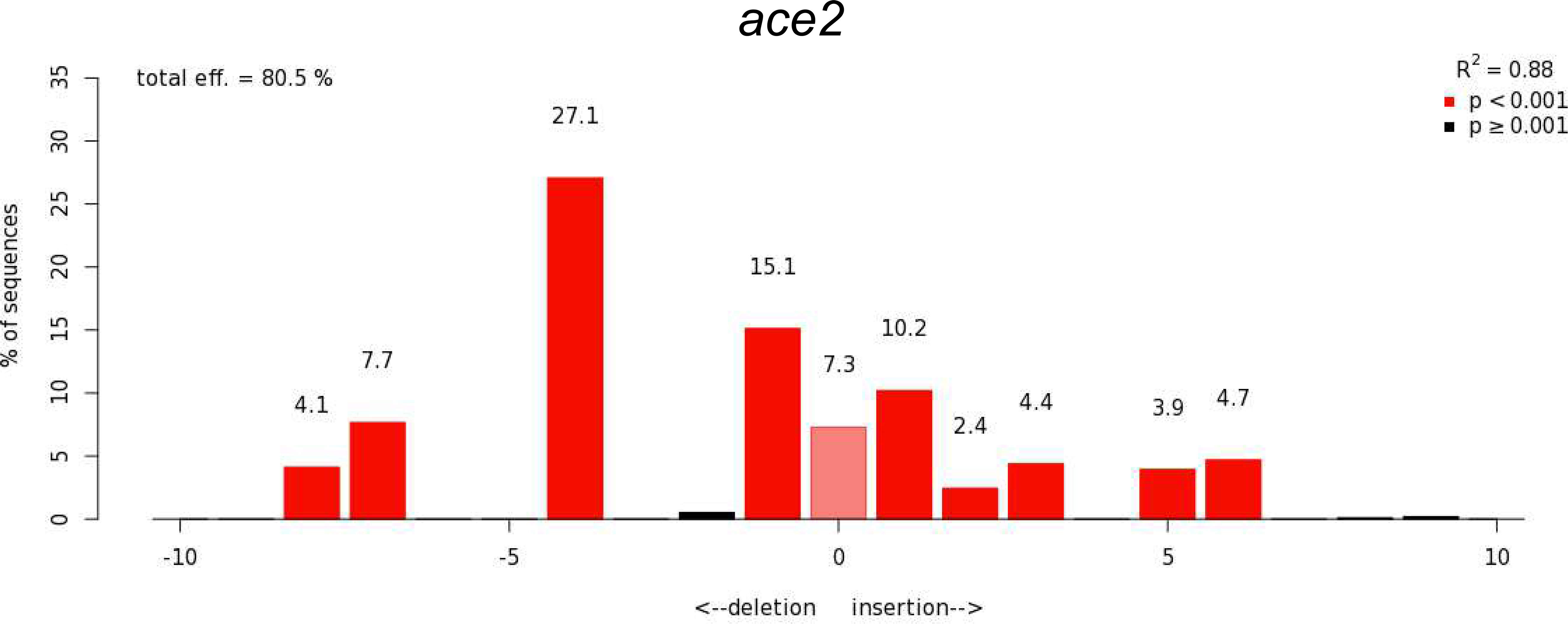
Efficiency of crRNA for ace2. Analysis of genome editing efficiency (80.5%) in larvae injected with ace2 crRNA/Cas9 complexes and quantification rate of nonhomologous end joining-mediated repair showing all insertions and deletions (INDELS) at the target site (https://tide.nki.nl/).

**Figure S5 (related to Figure 6).**
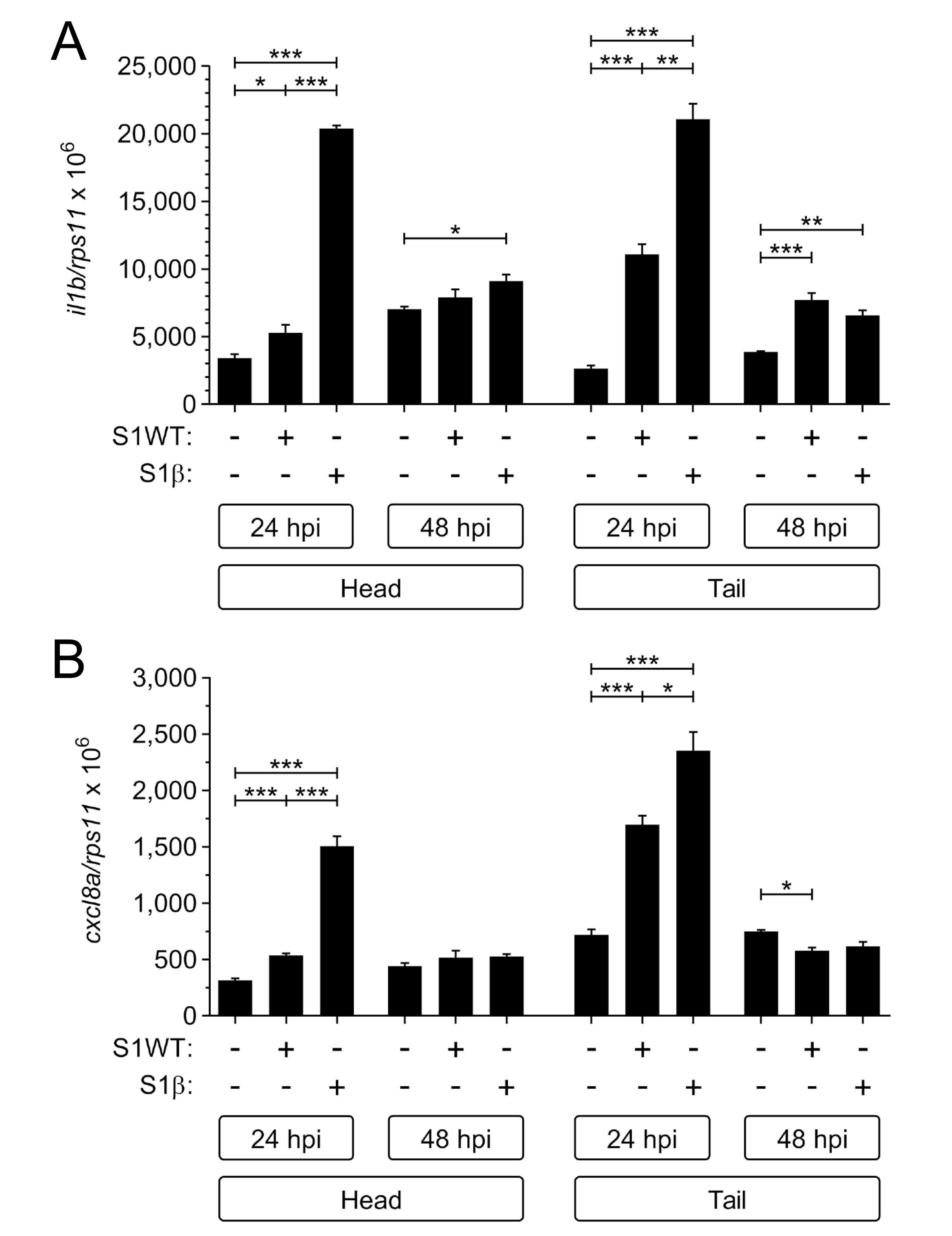
S1β shows delayed but stronger proinflammatory activity than S1WT. Recombinant S1WT or S1β were injected in the hindbrain ventricle (HBV) of 2 dpf wild type larvae. The transcript levels of *il1b* (A) and *cxcl8a* (B) were analyzed at 24 and 48 hpi by RT-qPCR. The data are shown as the mean ± S.E.M from 3 replicates. P values were calculated using one-way ANOVA and Tukey multiple range test. ns, not significant, *≤p0.05, **p≤0.01, ***p≤0.001.

**Table S1.**
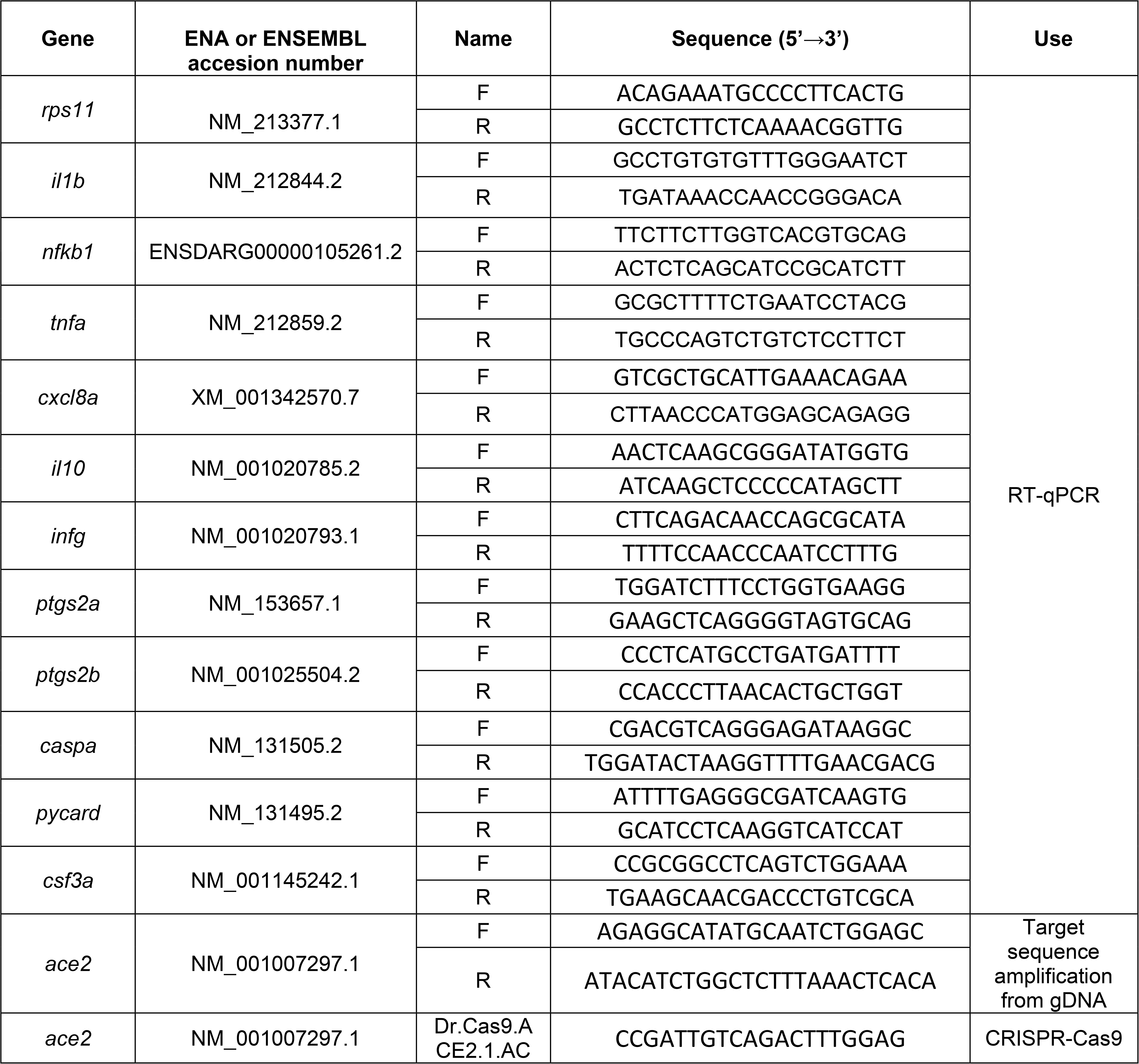
Primers and crRNAs used in this study. The gene symbols followed the Zebrafish Nomenclature Guidelines (http://zfin.org/zf_info/nomen.html). Ena, European Nucleotide Archive.

## Graphical abstract

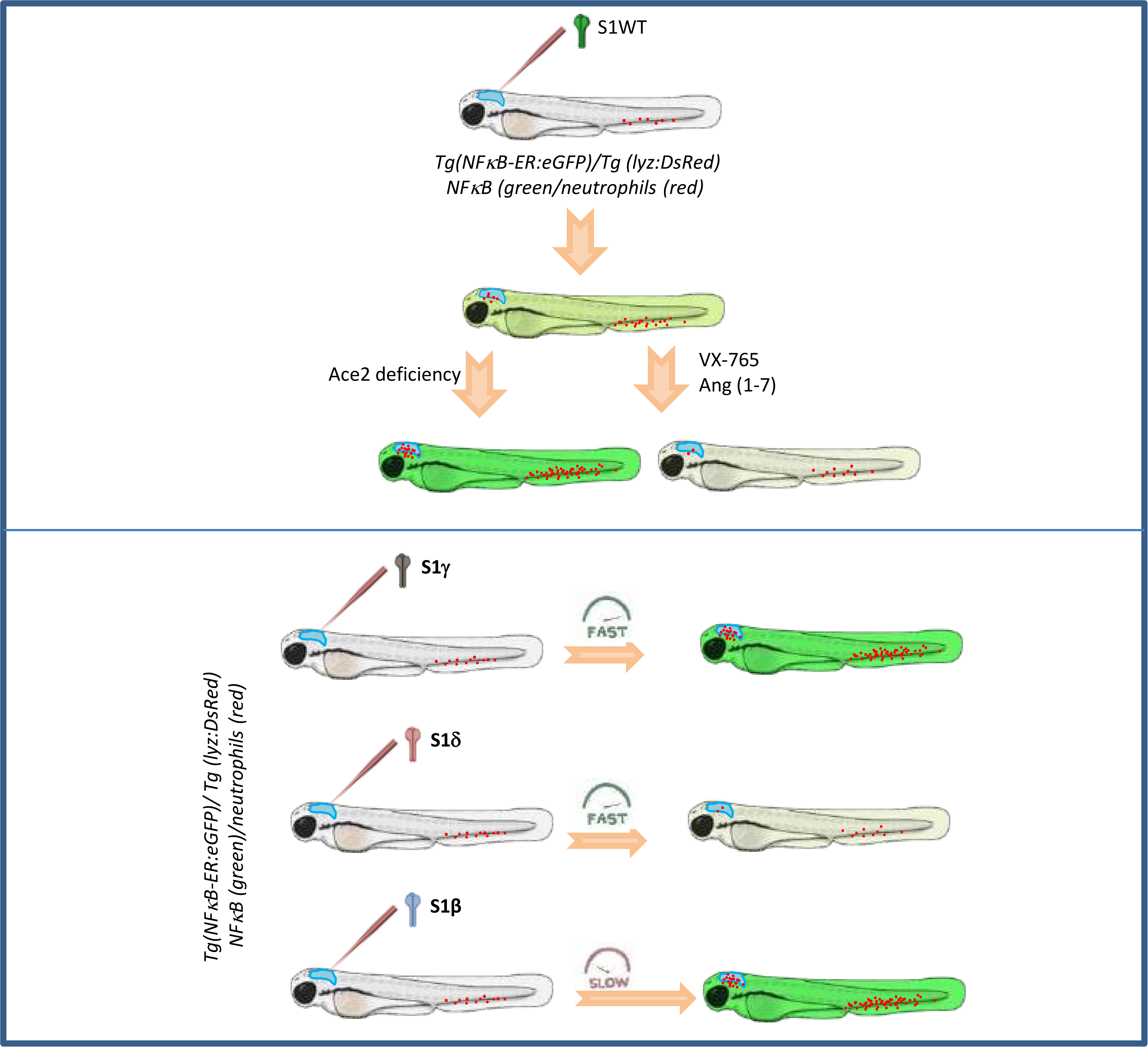

